# Shared species, strains and resistome between humans and pigs: A metagenomic analysis

**DOI:** 10.64898/2026.09.09.750220

**Authors:** Simeon Streit, Nadja Mostacci, Julia Moor, Anne Oppliger, Silas Kieser, Markus Hilty

## Abstract

Pig farming is one of the most intensive human–animal interfaces, and farm workers carry gut microbiotas that differ from those of non-farmers. Whether this overlap reflects genuine strain transmission or shared environmental exposure is unclear. Using shotgun metagenomics, we profiled the gut microbiomes of pigs at three growth stages (suckling, weaning, fattening), their farmers, and non-farmer controls, at species and strain resolution, and characterized the resistome in parallel.

Farmers shared more species with pigs than controls, with *Prevotellaceae* the most consistently enriched family. Strain-level analysis of 137 shared species showed that most (71%) maintain host-specific sub-species clades. Strain sharing ran an order of magnitude below species sharing and was confined to early growth stages; its direction could not be determined. Pigs and farmers shared a farm-associated resistome signature, including the beta-lactamase *cfxA4* and the methyltransferases *ermF* and *cfrE*. *ermF* and *tet(X)* were carried together on the same clonal Tn4351-family transposon, present in 86% of farmers and 80% of pigs but only 19% of controls. We found no evidence for transmission of mobile genetic elements between pigs and farmers; the shared resistome is best explained by carriage within shared gut taxa.

Species sharing, strain transmission, and resistance gene carriage thus reflect different scales of microbial exchange at the livestock–human interface. Our results argue against ongoing strain or mobile genetic element transmission from handled animals: the shared species and resistome are best explained by shared environmental exposure and by carriage of resistance genes within shared gut taxa.

## 1. Introduction

Pig farming is one of the most intensive human–animal interfaces globally (Pappas, 2013), and microbes can move between animals and their caretakers in either direction. Because humans and pigs are both omnivores with comparable gut physiology, a substantial overlap in gut microbial composition has been proposed (Liu et al., 2025); some have even proposed pigs as a better model than other animals for studying the human gut microbiota.

Intensive animal husbandry, and the associated use of antibiotics in livestock, is considered a key driver of the dissemination of antibiotic resistance genes (ARGs) into the environment and potentially into human gut microbiomes (Luiken et al., 2020). Farm workers spend long hours in barns and are repeatedly exposed to animal feces, airborne dust, and surfaces contaminated with ARG-carrying bacteria, which makes them a high-exposure group for resistance gene acquisition relative to the general population (Moor et al., 2021; Kraemer et al., 2019).

Occupational exposure to livestock environments has been associated with shifts in both the nasal and gut microbiota of farm workers. Studies of nasal microbiota in pig workers demonstrated that farm exposure is strongly linked to distinct microbial signatures reflecting the pig barn environment, with evidence for airborne animal-to-human bacterial transmission (Kraemer et al., 2019, 2018; Moor et al., 2021).

Moor et al. found that pig workers carried higher relative abundances of *Prevotellaceae* and lower abundances of *Bacteroidaceae* in their gut than controls (Moor et al., 2021; Sudatip et al., 2022). Amplicon sequence variants (ASVs) from *Prevotellaceae* were also detected simultaneously in pig feces, barn air, and pig worker stools, but not in cattle worker stools, supporting an airborne route of zoonotic microbial transfer (Moor et al., 2021). Consistent with this, *Prevotellaceae* are also enriched in the nasal microbiota of pig farmers compared to other farm workers, with airborne transmission risk reported to be highest in winter (Kraemer et al., 2019). Linked by the pharynx, microbes may be present in both the nose and oral cavity, which is relevant as oral–fecal transmission is a recognized process shaping the gastrointestinal microbiome in health and disease (Schmidt et al., 2019).

However, 16S rRNA gene sequencing is limited to genus-level resolution and cannot distinguish between species or strains, nor can it reliably detect antibiotic resistance genes (Johnson et al., 2019). Shotgun metagenomics overcomes these limitations, enabling simultaneous characterization of the microbiome at species and strain resolution, quantification of functional gene content including ARGs, and assembly of metagenome-assembled genomes (MAGs) (Liu et al., 2025).

A Europe-wide, cross-sectional study of 176 farms (Van Gompel et al., 2019b) found that macrolide and tetracycline use in pig farming was directly associated with increased macrolide resistance gene abundance, and that total antimicrobial use during fattening correlated with total ARG abundance. Antimicrobial treatment therefore shapes the resistome of the pig gut directly, which makes a joint analysis of the pig and human microbiome essential. Among the main food-animal species — cattle, chickens, and pigs — pig farming alone accounts for roughly 41% of antimicrobial use (Ardakani et al., 2024), and antimicrobial use in food animals has been quantitatively associated with antimicrobial resistance in humans (Emes et al., 2022; Ardakani et al., 2023). People occupationally exposed to livestock carry the consequences: pig slaughterhouse workers harbour higher abundances of tetracycline and macrolide resistance genes than the general population (Van Gompel et al., 2019a).

Here, we applied shotgun metagenomics to the same cohort of Swiss pig farms, profiling pigs at three growth stages (suckling, weaning, and fattening), pig farm workers, and a healthy non-farmer control group. We aimed to: (i) characterize shared species and strains between pigs and pig farmers; (ii) compare the resistome across exposure groups; and (iii) determine whether the observed species-level overlap is maintained by ongoing strain transmission.

## 2. Main

### 2.1. Highly similar microbiome composition by alpha and beta diversity

The study was based on a prospective cohort of 31 pig farms from Switzerland (Moor et al., 2021). Pigs were tracked individually by ear tag and sampled longitudinally at three growth stages (suckling, ∼2 weeks; weaning, 6 weeks; fattening, 16 weeks), totalling 2,437 rectal swabs. Individual swabs were pooled by farm and growth stage prior to shotgun metagenomic sequencing, yielding 92 pig pools. Stool samples were collected from 43 pig farm workers from the same farms, and 57 non-farmer controls from a public European cohort were included (Feng et al., 2015).

We analyzed the microbial community composition using MetaPhlAn v4.1, which relies on a database of species genome bins (SGBs). Each SGB represents a cluster of genomes with >95% average nucleotide identity (ANI).

Pigs had higher gut microbiome richness and diversity than humans: 794 observed species in pigs, versus 384 in farmers and 225 in controls (mean Shannon index 4.84, 3.88, and 3.58, respectively; Figure 1). Both metrics increased across growth stages (suckling → weaning → fattening). The dominant ordination axis separated humans from older pigs (weaning + fattening pigs; PERMANOVA R² = 0.2; Figure 1(c)), yet pig farmers clustered significantly closer to pigs than non-farmer controls (Cohen’s d = −3; Figure 1(d)). All three pig growth stages differed significantly by PERMANOVA, with suckling pigs more distinct from weaning and fattening than those two were from each other (Supplementary Table S1; pairwise PERMANOVA: Weaning vs Fattening R² = 0.14, p < 0.001).

**Figure 1.**
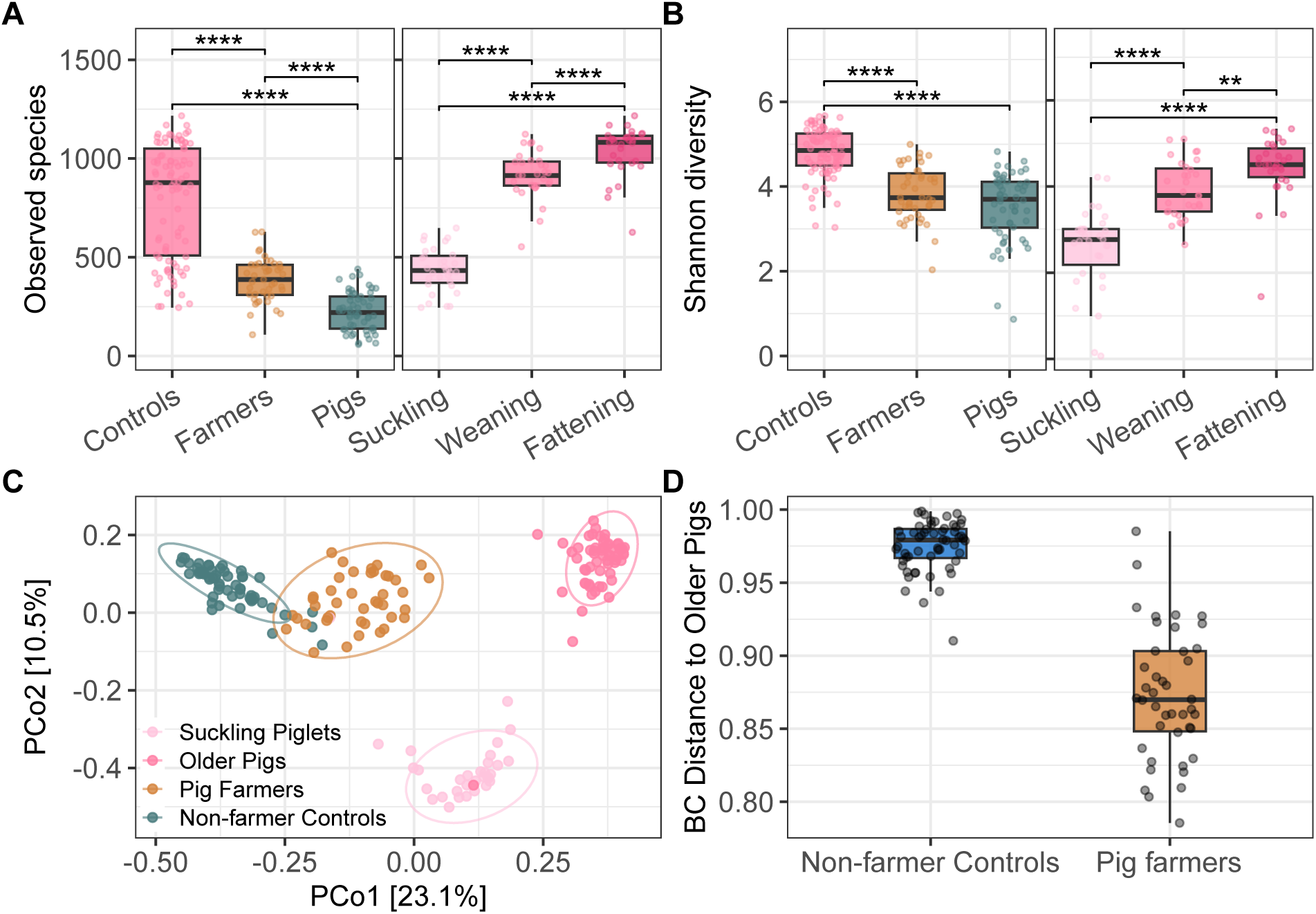
Microbiome *α*- and *β*-diversity. (A) Species richness and (B) Shannon diversity index per group. Significance tested using a linear model adjusted for total reads per sample. (C) Principal Coordinates Analysis (PCoA) on 3789 SGBs based on Bray-Curtis distance. The primary axis separates humans from older pigs (PERMANOVA R² = 0.2).(D) Pig farmers are closer to older pigs than non-farmer controls (Wilcoxon p = 5.5e-16, Cohen’s d = −3.022).

Farm identity was not a significant driver of beta diversity, as within-farm and between-farm Bray–Curtis distances were indistinguishable (Supplementary Figure S1). Weaning pigs were more similar to fattening pigs, even from other farms, than to suckling pigs from the same farm. Weaned pigs from different farms were often moved to a common fattening facility after weaning, which likely explains why the older-pig microbiome converges regardless of source farm.

#### 2.1.1. Shared species are enriched in Prevotellaceae

To quantify species-level overlap across groups, we assessed which SGBs were consistently detected in each group. We defined species presence as detection in 5% of samples with at least 10 reads and summarized group-level sharing using Venn diagrams (Figure 2(a)). Pigs harbored the largest number of unique SGBs. Farmers shared more species with pigs (653 SGBs) than with non-farmer controls (359 SGBs).

**Figure 2.**
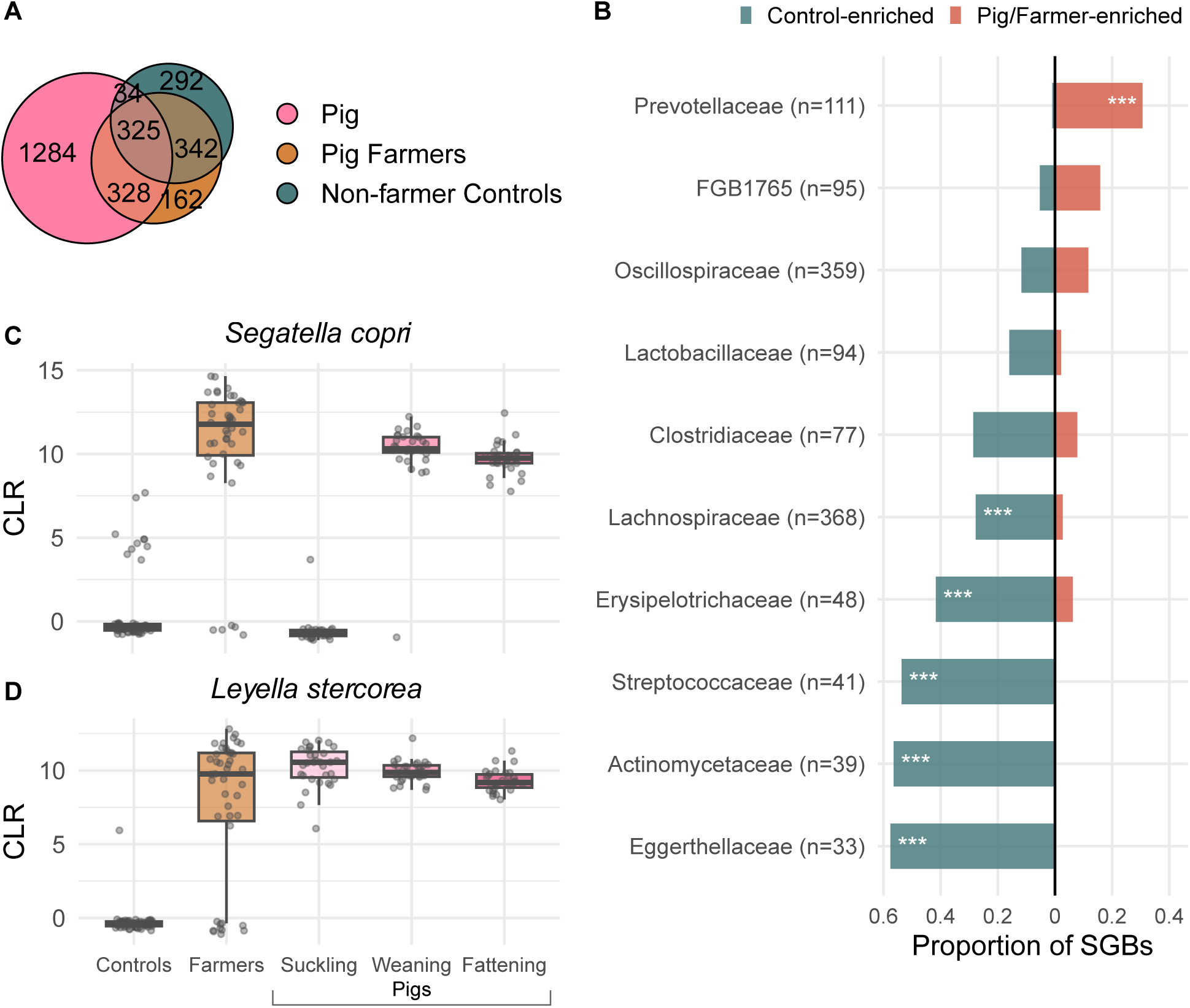
Shared species between humans and pigs. (A) Venn diagram showing the number of species genomic bins (SGBs) per group. (B) Proportion of differentially abundant SGBs per family. Red = enriched in pigs+farmers, teal = enriched in non-farmer controls. FGB1765 denotes a new *Firmicutes* family. Significance: Fisher exact test (BH-corrected). * p<0.05, ** p<0.01, *** p<0.001. Only older pigs included. (C) *Segatella copri* CLR abundance by group. (D) *Leyella stercorea* CLR abundance by group.

The species sharing rate between two samples (SGBs shared / SGBs in either sample) did not differ between same-farm and cross-farm comparisons in any group (Supplementary Figure S2).

Weaning and fattening pigs shared roughly half of their species with each other, while farmers shared one-eighth of their species with pigs, significantly more than non-farmer controls (5%).

To account for abundance in addition to presence/absence, we performed differential abundance analysis comparing farmers and pigs, separately, to controls (Supplementary Figure S3). Both comparisons identified *Prevotellaceae* as the most consistently enriched family (Figure 2(b)), matching prior 16S rRNA gene findings (Moor et al., 2021). Key representatives include *Segatella copri* (formerly *Prevotella copri*; Figure 2(c)), *Leyella stercorea* (Figure 2(d)), and *Prevotellamassilia timonensis*, also part of the *Prevotellaceae*. SGBs of *Lachnospiraceae* and *Actinomycetaceae*, in contrast, were predominantly depleted in pigs and farmers.

#### 2.1.2. A high proportion of shared species maintain host-specific or individualized lineages

To test whether species-level sharing reflects genuine exchange or merely shared taxonomy, we reconstructed strain-level phylogenies with StrainPhlAn for SGBs with sufficient marker coverage (n=137), and normalized the genomic distances (nGD) by the median tree distance to account for differences in evolutionary rates. For each SGB, we used multi-response permutation procedure (MRPP) on the normalized genomic distance matrix to test whether human and pig strains form distinct sub-species clades, and whether pig strains from the same farm are more similar than strains from different farms.

We saw that 71% (97/137) of SGBs formed distinct pig- and human-specific clades: most shared species therefore maintain separate evolutionary lineages in each host. *Bacteroidota* showed fewer host-specific clades than *Firmicutes* (Fisher’s exact test, p = 0.007; Figure 3(a)) and more frequent farm-specific clades (Fisher’s exact test, p < 0.001; Figure 3(b)), suggesting *Bacteroidota* species are more likely to cross the pig–human boundary, either directly or indirectly via unknown environmental sources.

**Figure 3.**
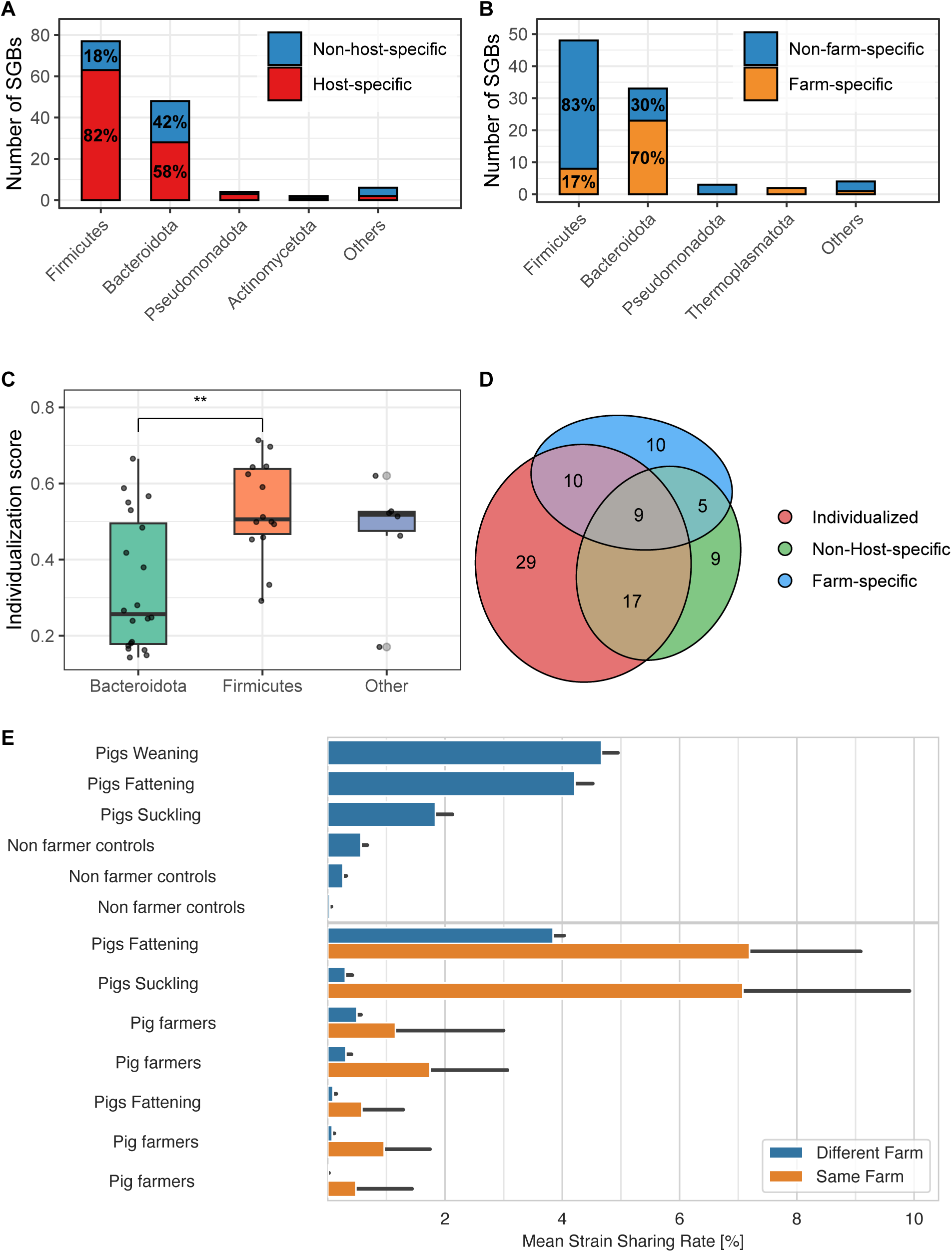
Sub-species host and farm specificity. (A) Number of host-specific and non-host-specific SGBs per phylum. (B) Number of farm-specific and non-farm-specific SGBs per phylum. *Bacteroidota* had fewer host-specific and more farm-specific SGBs than *Firmicutes*, consistent with more frequent cross-species exchange. (C) Individualization score by phylum. *Firmicutes* had more individualized SGBs than *Bacteroidota*, suggesting that strains from different individuals are more distinct in *Firmicutes*. (D) Venn diagram showing the overlap of SGBs classified as individualized, non-host-specific, and farm-specific. Only 14 SGBs were neither host-specific nor individualized. (E) Mean strain sharing rate between sample pairs by comparison type, split by same-farm vs. different-farm pairs.

An intermingled phylogenetic tree, in which human and pig strains are not separated into distinct clades, is often interpreted as evidence of species sharing. We considered an alternative explanation: the species may be highly individualized, with every individual carrying a distinct strain. To separate these two scenarios, we calculated an *individualization score* for each SGB (n = 137), defined as the median distance to the nearest neighbour averaged across all samples. *Firmicutes* had higher individualization scores than *Bacteroidota* (Figure 3(c)), meaning that strains of most *Firmicutes* species are more distinct between individuals, whereas *Bacteroidota* strains tend to be more similar across individuals and hosts. Most SGBs were either host-specific or individualized; 14 were neither (Figure 3(d)). The latter are the species whose strain structure is consistent with genuine *species* sharing between hosts.

#### 2.1.3. Strain sharing between pigs and pig farmers is rare

Strain-level resolution is the most stringent test of cross-host microbial exchange. We defined potential strain sharing as a normalized genomic distance (nGD) below 0.1, matching established StrainPhlAn cutoffs (Ferretti et al., 2018; Mostacci et al., 2023).

Surprisingly, the host-specific classification did not predict strain sharing. In fact, 72% of sharing events involved host-specific SGBs (Supplementary Table S6). *Escherichia coli* illustrates the point: it has a human-specific clade, yet other clades show strain transmission between humans and pigs (Supplementary Figure S4). In contrast, the individualization score was predictive: 88% of sharing events involved non-individualized SGBs, whereas only 12% involved individualized SGBs (Supplementary Table S6).

Strain sharing rates were an order of magnitude lower than species sharing rates: sharing was highest between weaning and fattening pigs (4%), followed by suckling pigs (1.8%), while pig farmers (0.51%) and controls (0.57%) both fell below 1% (Figure 3(e)). Unlike species sharing, strain sharing showed a strong farm effect: over 7% of strains persisted within farms across pig growth stages, whereas farmer–pig sharing was detectable on the same farm (0.86% with weaning pigs, 1.26% with suckling pigs) but near-absent across farms (0.08–0.31%). Similarly, control–pig sharing was virtually zero (0.03%).

The farmer–pig sharing events were confined to the early growth stages (Figure 3(e)). During the suckling stage, most events involved *Oscillospiraceae*; during the weaning stage, *Prevotellaceae* dominated (Supplementary Figure S5). For example, *Prevotellamassilia timonensis* had two strain-sharing events between farmers and weaning pigs on the same farm (Supplementary Figure S6).

With fattening pigs, we observed only five potential strain-sharing events in total (Supplementary Table S5), two of which came from farmers who had contact with the pigs only during the weaning stage, suggesting that the strains were acquired earlier and persisted into the fattening period.

### 2.2. The resistome mirrors taxonomic composition

Mapping reads to the CARD database, pigs and pig farmers carried more ARG genes than controls (mean 966 and 1,081 observed genes per sample, respectively, vs 878 in controls; Supplementary Figure S8), and ARG richness in pigs was highest in suckling piglets and decreased thereafter (suckling vs weaning: p = 0.019; suckling vs fattening: p = 0.001; weaning vs fattening: p = 0.57; linear model adjusted for sequencing depth, Tukey-adjusted pairwise comparisons; Supplementary Figure S9). Resistance genes belonging to tetracyclines were the most abundant resistance class across all groups, as expected given their ubiquity in gut metagenomes (Figure 4(a))(Wang et al., 2019; Chandel et al., 2024). MLS (macrolide, lincosamide, streptogramin) and beta-lactam ARGs were elevated in both pigs and pig farmers relative to controls (pairwise Wilcoxon rank-sum test, BH-adjusted: p < 0.001 for each comparison; Supplementary Figure S10), with no significant difference between these two groups. In contrast, log2 ratio analysis of ARG drug classes revealed that ARGs for aminoglycosides were significantly higher in pigs than either human group (BH-adjusted p < 0.0001), while mupirocin and glycopeptide ARGs (primarily human clinical use) were more abundant in non-farmer controls (Figure 4(b)). Suckling piglets showed the broadest ARG profile, likely because their immature gut microbiome is more receptive to colonization and exchange (Tang et al., 2025), and because antibiotic use in our data was also concentrated at this stage (Supplementary Table S3).

**Figure 4.**
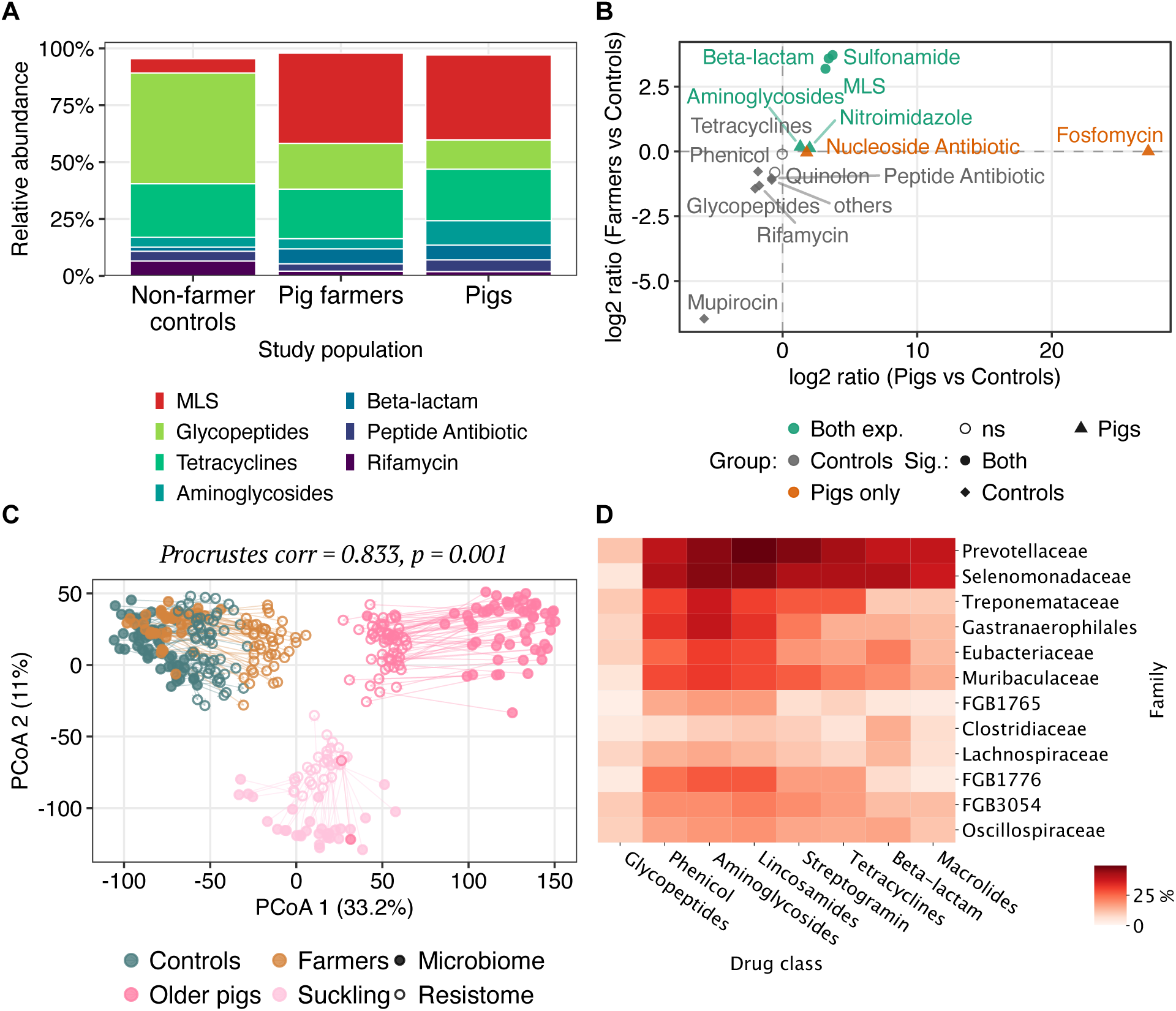
Resistome diversity and composition. (A) Relative abundance of ARG drug classes by group. (B) Log2 ratio of ARG drug classes between pigs, farmers, and controls. (C) Procrustes analysis aligning microbiome (filled circles) and resistome (open circles) CLR-based ordinations. Arrows connect each sample between the two spaces. Procrustes correlation = 0.833, p = 0.001. (D) Percentage of each family’s SGBs with at least one positive Spearman correlation ( > 0.5) to an ARG of the given drug class.

Beta-diversity of the resistome resolved into the same three clusters as the microbiome: humans (controls and farmers), suckling pigs, and older pigs (Figure 4(c)), and farmers were again significantly closer to pigs than controls in resistome space (Wilcoxon p < 0.001, Cohen’s d = 2.53; Supplementary Figure S12). Procrustes analysis confirmed a significant correlation between microbiome (SGB) and resistome (ARG) beta diversity (p = 0.001, correlation = 0.833; Supplementary Table S2), meaning taxonomic composition drives the overall resistome differences between groups. Consistent with this, the SGB–ARG correlations were dominated by *Prevotellaceae* and *Selenomonadaceae* (Figure 4(d)).

#### 2.2.1. Single ARGs correlate with Prevotellaceae species

To identify the individual ARG genes behind these drug-class patterns, we ran DESeq2 on all ARO-term-level ARG counts, testing pigs and farmers each against non-farmer controls. Genes that were significantly differentially abundant (FDR < 0.05, |log2FC| > 1) in the same direction in both contrasts were retained (Figure 5(a)). To guard against spurious read-mapping calls, genes were only retained if they were detected in 10% of samples.

**Figure 5.**
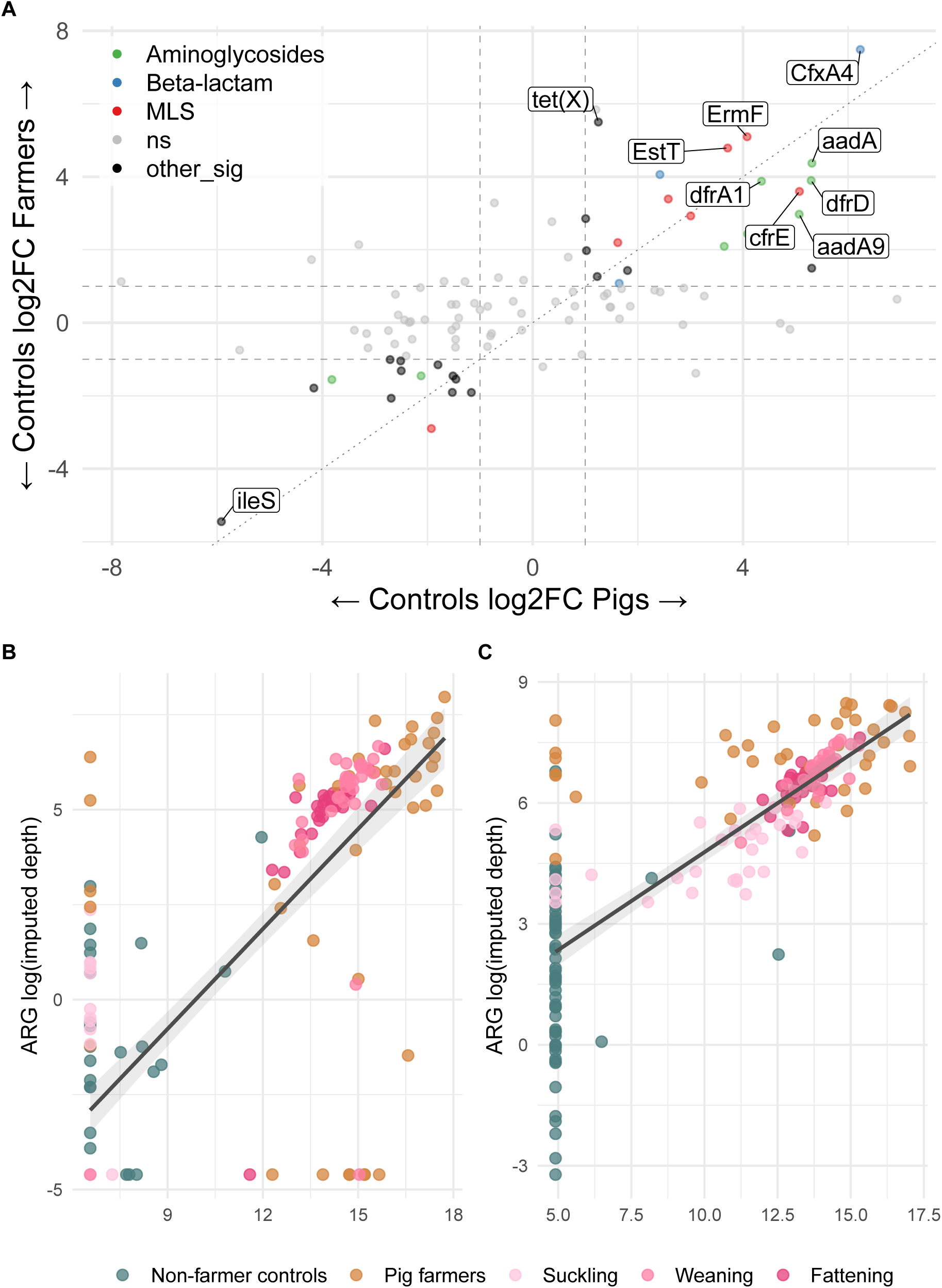
Differentially abundant ARGs and their *Prevotellaceae* hosts. (A) Log2 fold-change scatter of all ARGs (ARO terms, coloured by drug class) comparing pigs vs controls and farmers vs controls. (B) *cfxA4* vs *Segatella copri*: Spearman = 0.76, partial (adjusted for group) = 0.48. (C) *ermF* vs *Segatella sinensis*: Spearman = 0.79, partial (adjusted for group) = 0.42. Points are coloured by group; the grey line is the overall linear fit.

The ARGs most strongly elevated in both pigs and farmers include the beta-lactamase *cfxA4*, the 23S rRNA methyltransferases *ermF* and *cfrE*, and the tetracycline-inactivating monooxygenase *tet(X)* (Figure 5(a)). The most depleted ARGs included *ileS* (mupirocin resistance) and the glycopeptide resistance genes *vanU* and *vanY*, all significantly more abundant in non-farmer controls than in pigs or farmers (adjusted p < 0.001 in both contrasts), matching the drug-class pattern for mupirocin and glycopeptides.

We next asked whether the abundance of individual ARGs tracks that of individual species, using a correlation scan of all SGB–ARG pairs (n = 2,857,680) across pigs and farmers. Consistent with the family-level pattern above, the farm-associated ARGs were not tied to a single host: their strongest partners were multiple members of the *Prevotellaceae*, which accounted for 8 of the top 20 correlations for the beta-lactamase *cfxA4* and 7 of 20 for the 23S rRNA methyltransferase *ermF*. As representative examples, *cfxA4* was most strongly correlated with *Segatella copri* (Spearman = 0.76; Figure 5) and *ermF* with *Segatella sinensis* ( = 0.79), but each gene correlated similarly with several other *Prevotellaceae* species (e.g. *S. sinensis*, *S. brunsvicensis*, *Prevotella dentalis*). These correlations persisted after adjusting for group membership, indicating that the ARGs and their *Prevotellaceae* hosts co-vary beyond the group-level enrichment they have in common.

#### 2.2.2. ARG sharing in the absence of strain sharing

To test whether the farm-associated ARGs identified above can be mobilized, we annotated MGEs on all contigs carrying a CARD hit (MEFinder) and asked which ARGs share a contig with an MGE within 1 kb.

Of the 20 ARGs elevated in both pigs and farmers, MGE linkage (an MGE predicted within 1 kb on the same contig) was detected for six, but it was substantial for only two of these: *tet(X)* (79% of 81 loci) and *ermF* (46% of 94 loci), both carried on the same Tn4351-family transposon (Figure 6(a)). The remaining four showed low linkage rates (*dfrA1* 15%, *sul2* 14%, *estT* 11%, *aadA* 7%). *dfrA1* loci were associated with transposons and insertion sequences characteristic of *Enterobacteriaceae* plasmids (Tn2, Tn7, Tn6196, IS26, IS6100), consistent with integron-borne trimethoprim resistance on enterobacterial plasmids rather than the pig-gut-associated *Bacteroidetes* elements.

**Figure 6.**
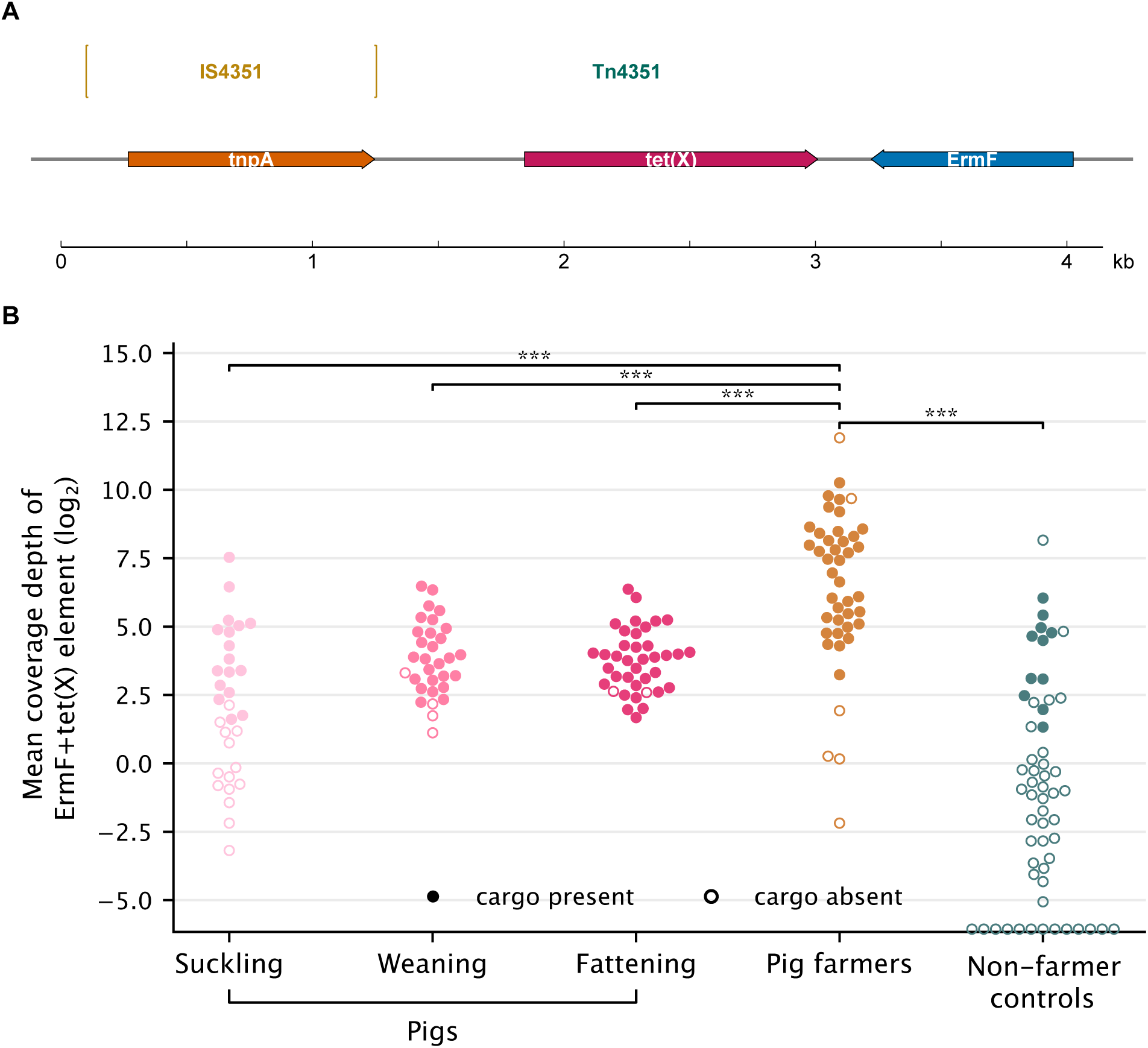
Farm-associated ARGs and their mobile genetic context. (A) Gene/domain architecture of the representative element (IS4351-flanked Tn4351 composite carrying *tet(X)* and *ermF*). Both *ermF* and *tet(X)* are carried on the same Tn4351-family transposon (45 contigs, 40 with a full MEFinder Tn4351 prediction; sequences ∼99.9% identical). (B) Mean per-base coverage depth of the *ermF*+*tet(X)*-carrying Tn4351 element per sample (log2 scale); filled circles denote samples passing the strict cargo test (whole element and each of *tet(X)*, *ermF* and Tn4351 covered 80 % at 1×), open circles denote absent/partial elements.

In 45 contigs, *ermF* and *tet(X)* were present on the same contig together with Tn4351, a *Bacteroides*-associated transposon on which *tet(X)* was first described (Speer et al., 1991). The 22 Tn4351 variant sequences were 99.7 % identical (median 99.95 %) across all 40 elements. Read mapping of the *ermF* +*tet(X)*-carrying Tn4351 element across all 200 samples showed that it was present (strict cargo test: whole element and each of *tet(X)*, *ermF* and the Tn4351 unit transposon covered 80 % at 1× depth) in 86% of farmers and 80% of pigs but only 19% of controls (Figure 6(b)). Farmers and pigs both carried the element significantly more often than controls (Fisher’s exact test, p < 0.0001 for each), whereas prevalence did not differ between pigs and farmers (p = 0.48). In pigs, presence increased across growth stages, from suckling (55%) through weaning (87%) to fattening (95%), the opposite of the strain-sharing pattern, which was confined to the early growth stages. Identical element sequence occurs in *Segatella copri* and *Bacteroides hominis* reference genomes, indicating carriage in multiple *Bacteroidetes* species.

Other farm-associated ARGs that were near-ubiquitous in the farm-exposed groups, such as *mecD*, *vanN*, *cfrE* and the *cfxA*-type beta-lactamases, showed essentially no MGE linkage ( 2% of loci; Supplementary Table S4).

## 3. Discussion

Building on the 16S rRNA gene study of Swiss pig farms by Moor et al. (2021), we used shotgun metagenomics to examine the gut microbiomes of pigs, their farmers, and non-farmer controls at the species and strain level, and measured the resistome in parallel. We found that humans and pigs shared many species, but strain transmission between them was rare.

### 3.1. Species sharing reflects occupational exposure, not shared hosts

Pigs harbor a highly diverse gut microbiome that increases with growth stage, with suckling piglets compositionally distinct from older animals. Pig farmers’ gut microbiome diversity was higher than in non-farmer controls but lower than in pigs, matching the prior 16S analysis (Moor et al., 2021). Farmers also clustered closer to pigs than controls in ordination space, confirming that occupational exposure shapes the human gut microbiome.

At the species level, farmers shared twice as many SGBs with pigs ( 11% species overlap) than non-farmer controls did ( 5%; Figure 2(a)). *Prevotellaceae* was the most consistently enriched family in both groups, matching the prior 16S study (Moor et al., 2021) and pig-farming cohorts from other countries (Sudatip et al., 2022).

### 3.2. Strain sharing is rare and confined to the early growth stages

Shared species do not necessarily indicate that a species was transmitted between pigs and humans. In 71% (97/137) of the shared species, human and pig strains formed distinct sub-species clades, indicating that most shared species maintain separate evolutionary lineages in each host.

An intermingled phylogenetic tree, in which human and pig strains are not separated into distinct clades, is often interpreted as evidence of sharing, but we showed that this is not the case. First, among the species with sufficient strain coverage, 47% were highly individualized. Second, our strain-level sharing analysis showed that intermingled phylogenies do not predict strain sharing.

Strain sharing rates were an order of magnitude lower than species sharing rates. Farmer–pig sharing was significantly higher within farms than across farms (0.86% with weaning and 1.26% with suckling pigs, compared with 0.08–0.31% across farms), whereas control–pig sharing was virtually zero (0.03%). As a reference, a baseline of <0.1% strain transmission is expected even between unrelated hosts (Heidrich et al., 2025). We therefore detect no ongoing transmission.

An earlier longitudinal study of dairy farmers similarly found that exposure effects on the gut community were only apparent at higher taxonomic and functional resolution (Mahmud et al., 2024). The pig gut microbiome is compositionally closer to the human gut than to the ruminant gut (Liu et al., 2025), yet compositional similarity does not predict the frequency of strain exchange.

Finally, our results pertain to the gut. The nasal and oral cavities, which are more directly exposed to barn air and dust and were previously shown to carry *Prevotellaceae* of likely porcine origin (Kraemer et al., 2019), may represent more permissive routes of livestock-to-human strain transmission than the mature gut ecosystem.

### 3.3. Why Prevotellaceae enrichment without ongoing strain transmission?

Our results argue against ongoing occupational transmission from the animals farmers currently handle, yet farmers and pigs share a high fraction of species, particularly *Prevotellaceae*. How, then, is this enrichment maintained?

Farmers may have acquired their strains from earlier contacts with previous pig cohorts; once established, these strains may resist displacement by new arrivals (priority effect). The widespread cross-farm strain sharing we observed further points to exchange through common environmental reservoirs (feed, water) and pig transport to fattening facilities, operating over longer timescales and broader networks than our cross-sectional snapshot captures. The true transmission rate may also be underestimated, as distinct metabolic and immunological niches (Tett et al., 2021) may drive rapid adaptation of transmitted strains.

The enrichment is not specific to pig farming: dairy farmers likewise show elevated gut *Prevotella* that scales with contact frequency with cattle (Cuperus et al., 2025), suggesting that elevated *Prevotella* is a general occupational signature across livestock systems. Whether this shared signature reflects a common mechanism (repeated low-level environmental exposure, or a general response to the farm environment) rather than livestock-specific strain transmission cannot be resolved with our cross-sectional data.

### 3.4. The shared resistome is carried within shared host taxa

Turning to the resistome, we found complementary patterns. At the community level the resistome mirrored the microbiome: beta-diversity resolved into the same group structure, and ARG composition correlated significantly with microbiome composition (Procrustes p = 0.001). Resistome differences are therefore largely microbiome-driven. The broader ARG carriage in pigs and farmers matches other European pig farm studies (Tams et al., 2023; Luiken et al., 2020), which likewise found no major shifts in farmers’ resistomes toward pig feces or dust resistomes (Luiken et al., 2020). Antimicrobial use in pig production plausibly drives these resistome patterns. A multinational metagenomic analysis of more than 1,000 gut metagenomes from humans and food animals found high abundance and diversity of ARGs in pigs and identified acquired ARGs shared between human and food-animal guts, with 345 ARGs shared between humans and pigs (Cao et al., 2022; Muhummed et al., 2025). In Switzerland, antimicrobial use in livestock is tracked through a nationwide electronic treatment journal: by weight, the highest amount of antibiotics is administered to fattening pigs and the lowest to suckling piglets, and although usage per pig (kg/pig) decreased between 2019 and 2021, treatment days and treatment incidence per farm increased (Wissmann et al., 2024). Measured as defined daily doses per animal (nDDD), however, suckling piglets and weaning pigs accounted for 50% and 44% of total antimicrobial consumption, respectively (Echtermann et al., 2020). This distinction between treatment amount and treatment frequency matters because early-life antimicrobial exposure shapes microbial composition and resistance gene acquisition (Vivarelli et al., 2026). Consistent with this, longitudinal shotgun metagenomic sampling of pig farm environments in low-antimicrobial-use Swedish production shows age-dependent increases in tetracycline and macrolide–lincosamide–streptogramin resistance that antimicrobial use alone cannot explain (Ladyhina et al., 2026). Despite regulatory progress under the Swiss Strategic Action Plan on Antibiotic Resistance (StAR) 2024–2027, pig farms thus remain a source of selective pressure for antimicrobial resistance, underscoring the importance of continued surveillance of resistance transmission routes in Switzerland.

The farm-associated ARG signature was nonetheless specific and coherent. Pigs and pig farmers carried more ARG genes than controls, with the same drug classes elevated in both groups: MLS (macrolide, lincosamide, streptogramin) and beta-lactam resistance, driven by the beta-lactamase *cfxA4*, the 23S rRNA methyltransferases *ermF* and *cfrE*, and the tetracycline-inactivating monooxygenase *tet(X)*. Aminoglycoside resistance was elevated specifically in pigs, while ARGs of primarily human clinical use (mupirocin, glycopeptides) were more abundant in controls. The signature was not tied to a single host: the farm-enriched genes co-varied specifically with the abundance of enriched *Prevotellaceae* species. The resistome signal is thus carried by the same taxonomic group that dominates species-level sharing between farmers and pigs.

The shared resistome is most parsimoniously explained by co-carriage within the shared gut taxa. The all-contig MGE screen showed that near-identical insertion sequences and transposons were broadly shared between pigs and farmers even though strain sharing was rare, with the same elements near-ubiquitous in both groups but largely absent from controls (Supplementary Figure S7). Because these elements belong to the *Bacteroidetes* taxa that dominate species-level sharing, and because they were present at no more than ∼1 copy per host genome, this pattern is best explained by genome-bound co-carriage rather than by independent element transfer; we therefore found no evidence that mobile genetic elements were transmitted between pigs and farmers. The most relevant element is the *tet(X)*/*ermF* -carrying Tn4351 transposon: a composite *Bacteroidetes*-associated element whose resistance cargo is flanked by two directly repeated IS4351 copies (Rasmussen et al., 1986; Mahillon and Chandler, 1998; Speer et al., 1991; Rasmussen et al., 1987), which was most prevalent precisely where strain sharing was absent: it rose across pig growth stages to 95% of fattening pigs and was present in 86% of farmers.

The species-spanning distribution of this element ( 99.7% identical across independent assemblies from many individuals and farms, and present in *Phocaeicola coprophilus* and *Leyella lascolaii* among our element-carrying contigs and in *Segatella copri* and *Bacteroides hominis* reference genomes) shows that it has transferred between *Bacteroidetes* species at some point; strain transmission, which operates within a single species, cannot account for this distribution. The resistance genes therefore retain the potential for gene-level dissemination, which is relevant for clinical antibiotic resistance risk. However, whether the element’s presence across the pig–farmer continuum reflects horizontal transfer or co-carriage within the shared host taxa cannot be established from these cross-sectional, short-read data: the element was present at no more than ∼1 copy per host genome and was consistently recovered integrated in contigs, consistent with genome-bound carriage. Resolving the two scenarios would require longitudinal sampling with matched time points and long-read sequencing to link elements to their hosts.

These findings should not be over-interpreted: only a fraction of loci were detectably MGE-linked, short-read assemblies capture such linkage only when ARG and MGE co-assemble onto the same contig, and co-occurrence does not demonstrate de novo transfer; establishing genuine MGE association would require long-read sequencing.

### 3.5. Limitations

Pig samples were collected longitudinally but pooled per farm and growth stage for sequencing, so per-animal strain tracking was not possible; within-farm strain persistence across stages therefore combines within-host persistence and transmission between littermates. Human samples were cross-sectional, preventing causal inference and definitive determination of transmission direction; the direction at the suckling stage remains uncertain, as sharing could reflect farmer-to-piglet transmission, piglet-to-farmer transmission, or a common environmental source, and longitudinal matched sampling would be needed to resolve this. StrainPhlAn analyses were restricted to SGBs with sufficient coverage ( 5 markers in 5 samples per group), potentially under-representing low-abundance taxa, and the strain-sharing analysis is cut-off-based: the 0.1 nGD threshold may miss true sharing events or include false positives.

ARG analysis captures gene presence, not expression or chromosomal mutations. ARG–MGE linkage was assessed on assembled contigs, where short contigs may break up genuine linkage; the reported linkage proportions are therefore conservative lower bounds, and long-read sequencing would be needed to confirm true MGE association. The all-contig MGE screen is read-based, with presence called at 50% element coverage, so it may miss or over-call partial elements; it cannot distinguish direct exchange from a common environmental source. Short-read metagenomes likewise cannot prove that the shared elements are genome-bound passengers rather than independently mobile: the elements were consistently recovered integrated in contigs and at no more than ∼1 copy per host genome, which is consistent with genome-bound carriage, but definitively resolving element–host linkage requires long-read sequencing. In addition, the 1,989 reference elements were clustered from pig and farmer assemblies only, and controls were screened by read mapping against this pig/farmer-derived reference; control-specific elements are therefore not represented, and the farmer/control prevalence comparisons concern elements that are at least present in the pig–farmer continuum rather than the full MGE repertoire of either group. Dietary intake was not profiled, so we cannot disentangle occupational exposure from diet-driven microbiome differences. Finally, our control samples were not processed identically to our farm samples but derived from previously published data; the similar ARG carriage observed in the Dutch pork production chain (Van Gompel et al., 2020) supports the validity of our approach.

### 3.6. Conclusion

We examined the pig–human gut microbiome interface at species, strain, and resistome resolution across multiple farms and pig growth stages. The strong species-level overlap between pig farmers and their animals, particularly the *Prevotellaceae* enrichment reported across livestock systems worldwide, was not matched by correspondingly high strain transmission. Most shared species either maintain host-specific lineages or are highly individualized, and the few strain-sharing events we detected were confined to early growth stages, their apparent rate inflated by long-lived strain persistence within the pig pool. Species-level overlap therefore most likely reflects shared environmental exposures operating over longer timescales and strain acquisitions during earlier contact with previous pig cohorts, rather than ongoing transmission from currently handled animals.

Resistome differences between groups were primarily driven by the underlying microbiome composition. The farm-enriched ARGs *ermF* and *tet(X)* are carried on a Tn4351-family transposon enriched across the pig–farmer continuum; its occurrence across multiple *Bacteroidetes* species leaves these genes with the potential for gene-level dissemination. We nonetheless found no evidence that this element, or other mobile genetic elements, moved between pigs and farmers, and the shared resistome is best explained by carriage within the shared host taxa.

Species sharing, strain transmission, and resistance gene carriage thus reflect different scales of microbial exchange at the livestock–human interface. Only species-level overlap showed evidence of ongoing exchange between pigs and farmers.

## 4. Methods

### Samples and sequencing

Samples were obtained from a previously established cohort of Swiss pig farms (Moor et al., 2021). Briefly, prospective sampling was performed on pig breeding farms in Switzerland between March 2018 and March 2019 in the cantons of Vaud, Bern, and Fribourg. Pigs were tracked individually by ear tag and sampled longitudinally at three life stages (suckling, ∼2 weeks old; weaning, 6 weeks old; fattening, 16 weeks old), totalling 2,437 rectal swabs (918 suckling, 866 weaning, 653 fattening) from 31 farms. Pig farm workers (n=43) from the same 31 farms self-collected stool samples. Ethical approval was obtained from the Human Research Ethics Committee of the Canton of Vaud (2018-00080) and the Veterinary Ethics Committee of the Canton of Vaud (VD3335). DNA was extracted using the QIAamp DNA Mini Kit following the manufacturer’s instructions as described in (Moor et al., 2021). In contrast to the original study, samples were pooled by farm and life stage prior to shotgun metagenomic sequencing (Illumina NovaSeq 6000, PE150), generating ∼10 Gb of raw data per sample. This yielded 92 pig pools (31 suckling, 31 weaning, and 30 fattening), each representing one farm at one growth stage. Together with 43 pig farm workers and 57 public human control metagenomes from a non-exposed European population (ENA accession ERP008729, Feng et al. (2015)), the dataset comprised 192 unique metagenomes; eight additional libraries were technical duplicates of pig pools and were excluded from the differential abundance analyses.

### Read processing and taxonomic profiling

Adapter removal, quality filtering and deduplication were performed with fastp (v0.23.4) (Chen, 2023). Host (human and pig) reads were removed using BBMap (v39.10) against GRCh38 and Sscrofa11.1 references (Bushnell, 2014). Taxonomic profiling was performed with MetaPhlAn4 (v4.1.1) using the mpa_vJan25_CHOCOPhlAnSGB_202503 database to obtain species-bin-level relative abundances and estimated read mapping to full genomes (Blanco-Míguez et al., 2023). Analyses were run on the UBELIX HPC (University of Bern) for compute-heavy steps.

### Core microbiome and prevalence analysis

Species prevalence was calculated within each group (pigs, pig farmers, controls) using a 5% threshold. Overlap of species among groups was visualized using Venn diagrams (eulerr package, v7.0.4). Shared species-level genome bins (SGBs) between pigs and pig farmers were retained for downstream strain-level profiling.

### Strain-level analysis

Strain-level profiling was performed with StrainPhlAn4 (v4.1.1) using MetaPhlAn SAM outputs to extract consensus marker sequences; markers present in 50% of samples and samples with 5 markers were retained for phylogenetic reconstruction (Truong et al., 2017, Blanco-Míguez et al. (2023)). Resulting tree-based genomic distances were normalized by median tree length to obtain normalized genomic distances (nGD). Strain sharing was defined as an nGD below 0.1, and host-specificity and individualization were evaluated using Multi-Response Permutation Procedures (MRPP) and nearest-neighbour distance statistics.

Host-specificity was classified using MRPP on normalized genomic distance matrices (Euclidean distance, 999 permutations), with SGBs considered host-specific if within-group distances were significantly smaller than between-group distances (p < 0.05). Farm-specificity was tested analogously, comparing strains from the same farm against strains from other farms. The individualization score, defined as the median distance to the nearest neighbour averaged across all samples, quantified how distinct strains are between individuals; SGBs with high scores were classified as individualized. The full classification scheme is described in the supplementary material.

### Statistical analyses

Downstream analyses were performed in R (v4.1.1) (R Core Team, 2024) using packages including phyloseq (v1.50.0), DESeq2 (v1.46.0), vegan (v.2.7.2), and tidyverse (v2.0.0). *α*-diversity (Shannon, Simpson), richness, and Pielou’s evenness were calculated on species-level and ARO-aggregated CPM matrices. *α* -metrics were corrected for sequencing depth using a linear model. Post-hoc pairwise comparisons were performed with Tukey-adjusted estimated marginal means. *β*-diversity was assessed with Bray–Curtis dissimilarities (PCoA) and tested by PERMANOVA (adonis2, 999 permutations). Differential abundance for taxa and ARGs was performed with DESeq2 (v.1.46.0) on count data (taxa and ARGs with <10 reads were filtered), reporting features with |log2FC| > 1 and adjusted p 0.05 (Love et al., 2014).

DESeq2 was run with a design formula of ∼ category, where category encoded the group membership (pig, pig farmer, control). Both contrasts (pigs vs controls, farmers vs controls) were extracted from the same model. Independent filtering was applied, and log2 fold-change shrinkage was performed using the apeglm method (Salim et al., 2019).

For ARGs, reads were only counted for genes whose mapped reads covered 60% of the reference sequence in 10% of samples (RGI bwt Percent Coverage, max across alleles per ARO term), and counts were analysed both at the ARO-term level and after summing member genes within each AMR gene family. This coverage filter was applied once at the count/CPM matrix stage and thereby affected all resistome analyses (richness and diversity, drug-class composition, beta-diversity, Procrustes, and SGB–ARG correlations).

### Resistome annotation

Antibiotic-resistance genes (ARGs) were identified using RGI (CARD database, WildCARD/Resistome v4.0.1) and ARO term annotations. Chromosomal point mutations and broad efflux-pump annotations were excluded prior to the analysis (Alcock et al., 2022). Raw ARG counts were length-normalized by effective gene length

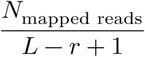

and converted to counts-per-million (CPM) for relative-abundance analyses. Low-abundance CPM (<1) values were set to zero.

### Drug class analysis

ARG abundances were aggregated to ARO terms and then harmonized into drug classes following the scheme of Munk et al. (2018) (Munk et al., 2018). When an ARO term mapped to multiple drug classes, abundance was split equally among classes (weight = 1/*n*). CPM values were log-transformed as *log*_10_(*CP M* + 10^−6^) for visualization and non-parametric testing. Group differences in drug-class abundance were assessed with pairwise Wilcoxon rank-sum tests (Benjamini-Hochberg corrected, *p_adj_* ≤ 0.05). Effect sizes were quantified as log median CPM ratios between groups.

### Contig assembly and ARG genomic context

Host-filtered, quality-trimmed reads were assembled per sample with SPAdes v4.2.0 (Nurk et al., 2017). ARGs were annotated on the resulting contigs with RGI (CARD database, --include_loose), and for each ARG hit the coding sequence (CDS) was extracted for downstream analysis. Taxonomy was assigned to each contig by best hit against reference genomes.

### Mobile genetic element (MGE) analysis of farm-associated ARGs

Contigs from all samples were screened for ARGs with RGI (CARD database, --include_loose), and MGEs were predicted on all ARG-carrying contigs with MEFinder 1.1.2. An ARG was classified as MGE-associated if an MGE was predicted on the same contig within 1 kb of the ARG. ARGs were retained for this analysis if they were significantly increased (log2 fold change and adjusted p-value, coverage-based) in both pigs and farmers relative to controls and had coverage 1× in at least one group (22 ARGs). To test whether the two farm-associated ARGs form a single horizontally transmissible element, contigs carrying both *ermF* and *tet(X)* were selected (n = 45) and a contiguous carriage element spanning Tn4351 + *tet(X)* + *ermF* was extracted from each (2.17–2.84 kb). All 45 elements were 99.7 % identical and co-linear, so a single 2,768 bp consensus reference was built by aligning them with minimap2 (-ax asm5) and calling the majority base; ORFs were verified by translation (*tet(X)* 388 aa, forward; *ermF* 267 aa, reverse). Reads from all 200 samples were mapped to this consensus with minimap2 (-ax sr --eqx --secondary=no -N 1), and per-base depth was computed with samtools depth -a. Reads were retained only if 95 % identical to the element (NM from the --eqx output), excluding divergent IS4351/transposase homologs. A sample was called **cargo-present** only if the whole element and each of its three components (the Tn4351 unit transposon, *tet(X)*, and *ermF* ) were each covered 80 % of their length at depth 1×; element abundance was reported as mean per-base coverage depth. Presence rates were compared between groups with Fisher’s exact test. Host taxonomy of element-carrying contigs was assigned with sendsketch (BBMap, RefSeq sketch database). Most contigs were too short to classify beyond *Bacteroidetes*; among the few with sufficient flanking sequence, element-carrying contigs were classified as *Phocaeicola coprophilus* and *Leyella lascolaii*. The identical element sequence was additionally identified in *Segatella copri* and *Bacteroides hominis* reference genomes by BLAST search against NCBI.

### All-contig MGE screen

To test whether MGE sharing extends beyond ARG-linked elements, MEFinder 1.1.2 was run on every assembled contig of the 143 pig and farmer samples, and all predicted element sequences were clustered within MGE families at 95% nucleotide identity over 80% of the shorter sequence, yielding 1,989 nucleotide-level elements. Reads from all 200 samples were mapped to the representative element sequences with BBMap (minid = 0.95), and an element was called present in a sample if 50% of its length was covered by mapped reads. Per-element prevalence was compared between pig farmers and non-farmer controls with Fisher’s exact tests and Benjamini-Hochberg FDR adjustment.

### SGB–ARG correlation analysis

To test whether specific taxa carry the farm-associated ARGs, we computed Spearman correlations between log1p-transformed SGB estimated counts (MetaPhlAn4) and log1p-transformed ARG depth (read mapping to the gene catalog) across all samples, focusing on *Prevotellaceae* SGBs and focal ARGs (*ermF*, *cfxA4*, *cfxA2*, *IMP-63*, *lnuC*, *tet(X)*, *tet*(O/W/Q)). Correlations were computed on samples where both the SGB and the ARG were detected ( 10 samples). To ensure that correlations were not driven merely by group-level co-enrichment, partial Spearman correlations were computed after regressing out group membership (category_with_pigstage) from the ranked values. An SGB–ARG pair was considered double-enriched if both the SGB (log2 fold change) and the ARG (log2 fold change) were increased in pigs/farmers relative to controls (log2FC > 1).

### Procrustes analysis

To assess concordance between microbiome and resistome composition, both SGB (MetaPhlAn4) and ARO-term (CARD) abundance matrices were CLR-transformed and subjected to Procrustes analysis using the protest() function from the vegan R package (999 permutations). The correlation coefficient R reflects the degree of configuration concordance between the two compositional spaces.

## 6. Post matter

### 6.1. Ethical approval

Ethical approval for this study was given by the Human Research Ethics Committee of the Canton of Vaud (2018-00080) and the Veterinary Ethics Committee of the Canton of Vaud (VD3335).

### 6.2. Funding

This study was supported by the Swiss National Science Foundation (Antimicrobial Resistance NRP72; research grant No. 177452 to M.H. and A.O.).

### 6.3. Data availability

The shotgun metagenomic sequencing data generated in this study are deposited in the Sequence read archive (SRA) under accession PRJNA1510222.

### 6.4. Acknowledgements

We thank the pig farmers and study participants for providing samples, and the staff of the Institute of Infectious Diseases, University of Bern, for technical assistance.

### 6.5. Author contributions

- **Conceptualization:** M.H., A.O.
- **Methodology:** N.M., S.K.
- **Software:** S.S., N.M., S.K.
- **Investigation:** S.S., S.K.
- **Resources:** J.M., A.O.
- **Data curation:** S.S., N.M.
- **Writing – original draft:** S.S., S.K.
- **Writing – review & editing:** S.S., S.K., M.H.
- **Visualization:** S.S., S.K.
- **Supervision:** S.K., M.H.
- Project administration: M.H.
- **Funding acquisition:** M.H., A.O.

### 6.6. Declaration of generative AI and AI-assisted technologies

During the preparation of this work, the authors used generative artificial intelligence (AI) tools in a supervised manner to assist with (i) writing and debugging the statistical-analysis code, (ii) searching literature and brainstorming interpretations, and (iii) editing and polishing the text of the manuscript. All AI-generated output was reviewed and validated by the authors, who take full responsibility for the content of this publication. No generative AI tool was used to generate or alter reported data, figures or images.

## 7. Supplementary

### 7.1. Supplementary Tables

### 7.2. Microbiome Diversity

### 7.3. Species Sharing

### 7.4. Strain-Level Analysis

### 7.5. Mobile Genetic Elements

### 7.6. AR Genes

**Supplementary Figure S1:**
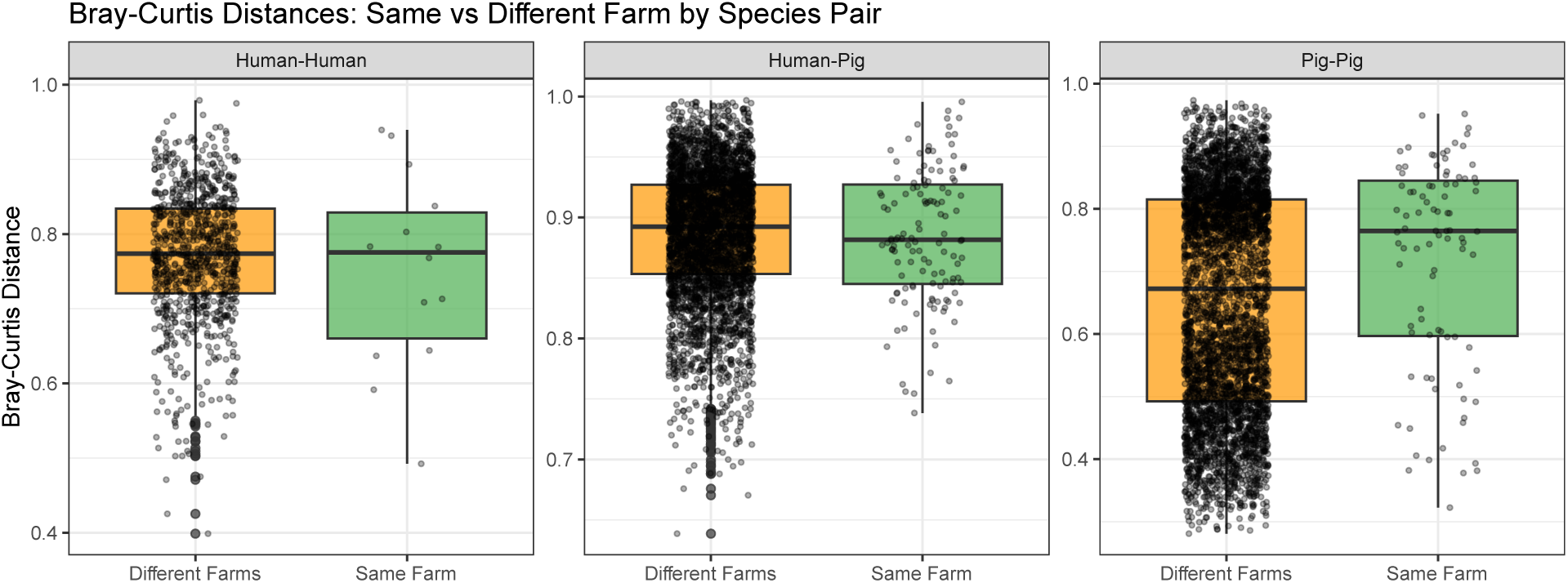
Beta diversity within vs. between farms.

**Supplementary Figure S2:**
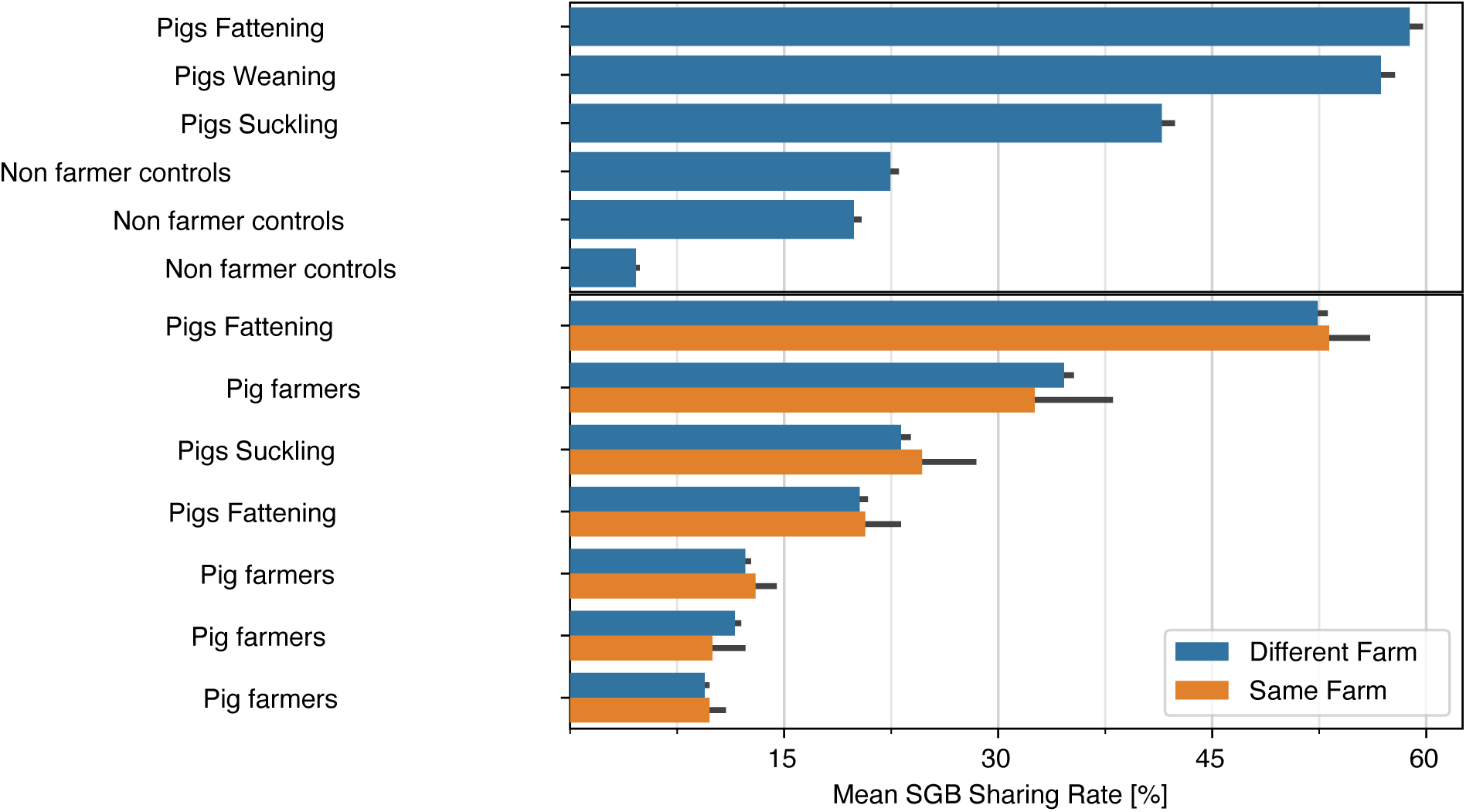
Species sharing rate between sample pairs (shared SGBs / total SGBs in either sample).

**Supplementary Figure S3:**
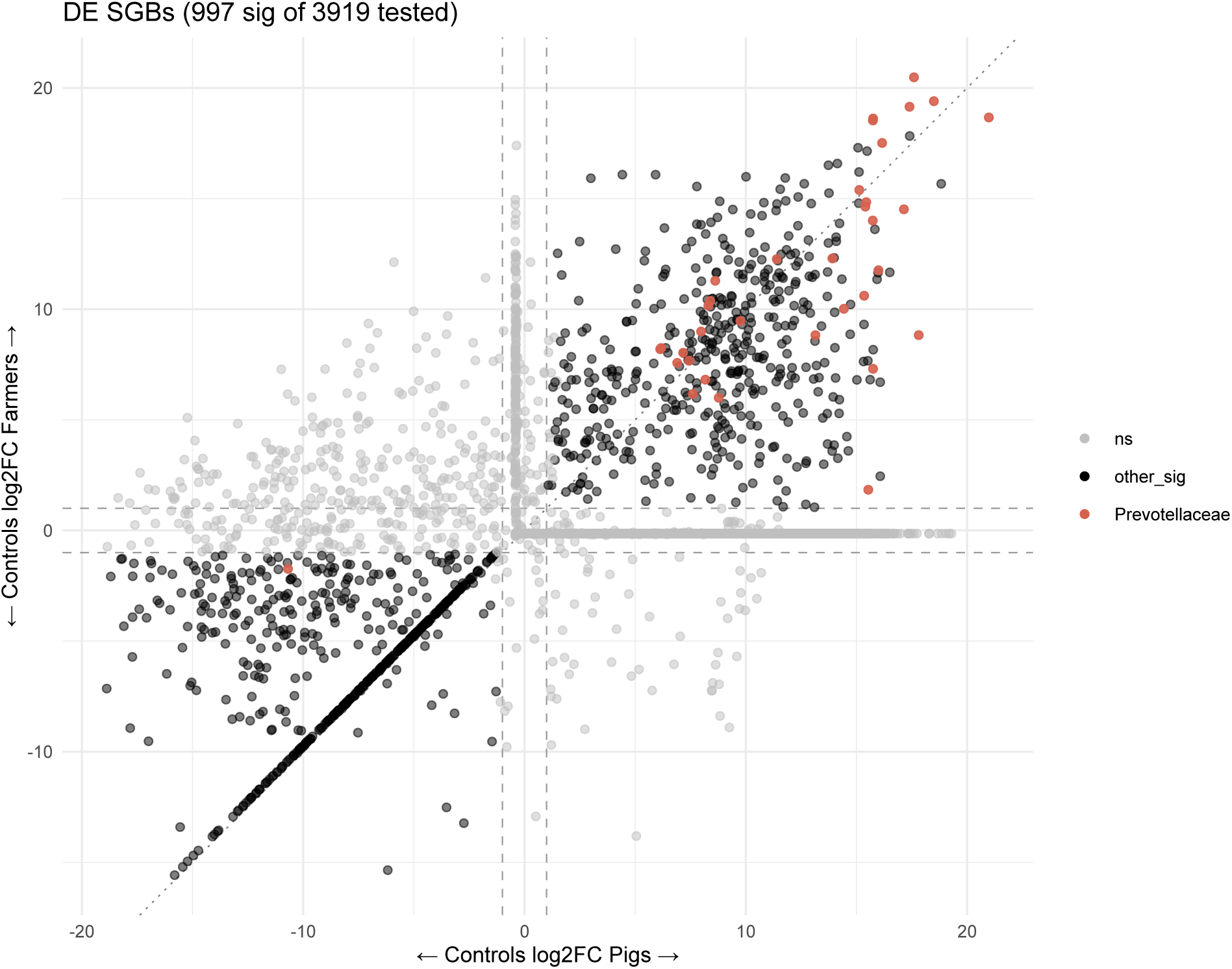
log2 fold-change scatter of all SGBs (Prevotellaceae highlighted).

**Supplementary Figure S4:**
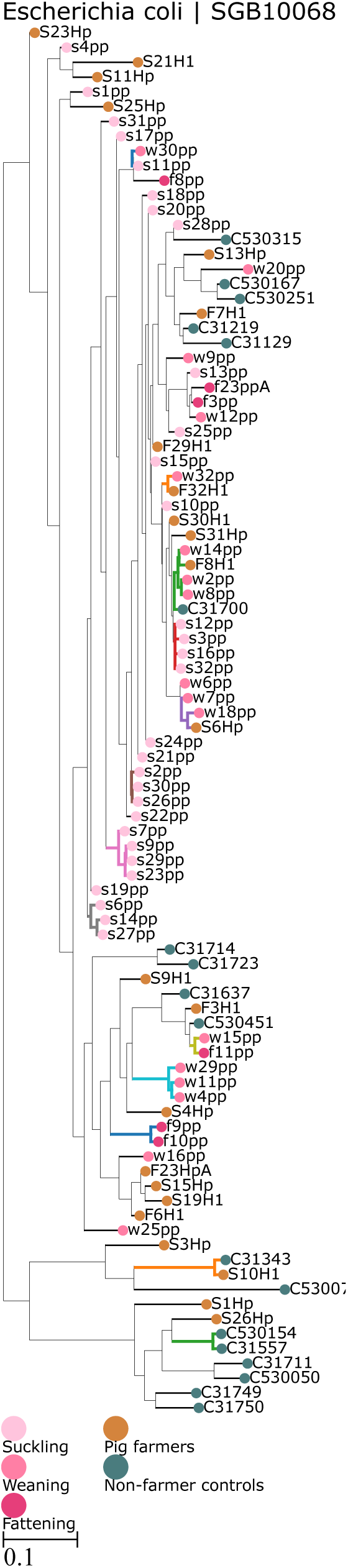
Phylogenetic tree of *Escherichia coli* strains (SGB10068).

**Supplementary Figure S5:**
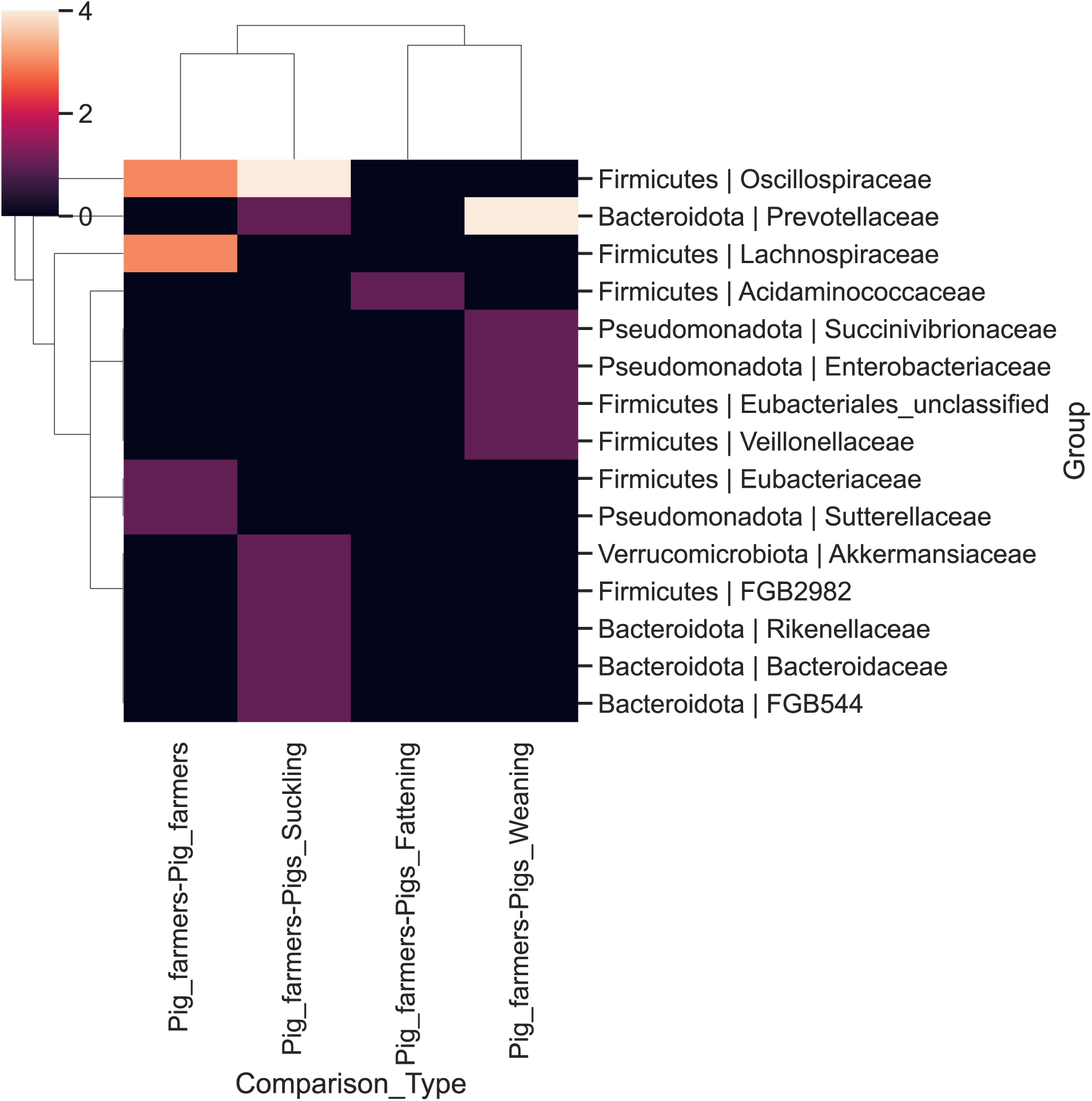
Number of shared strains with pig farmers on the same farm, aggregated by taxonomy.

**Supplementary Figure S6:**
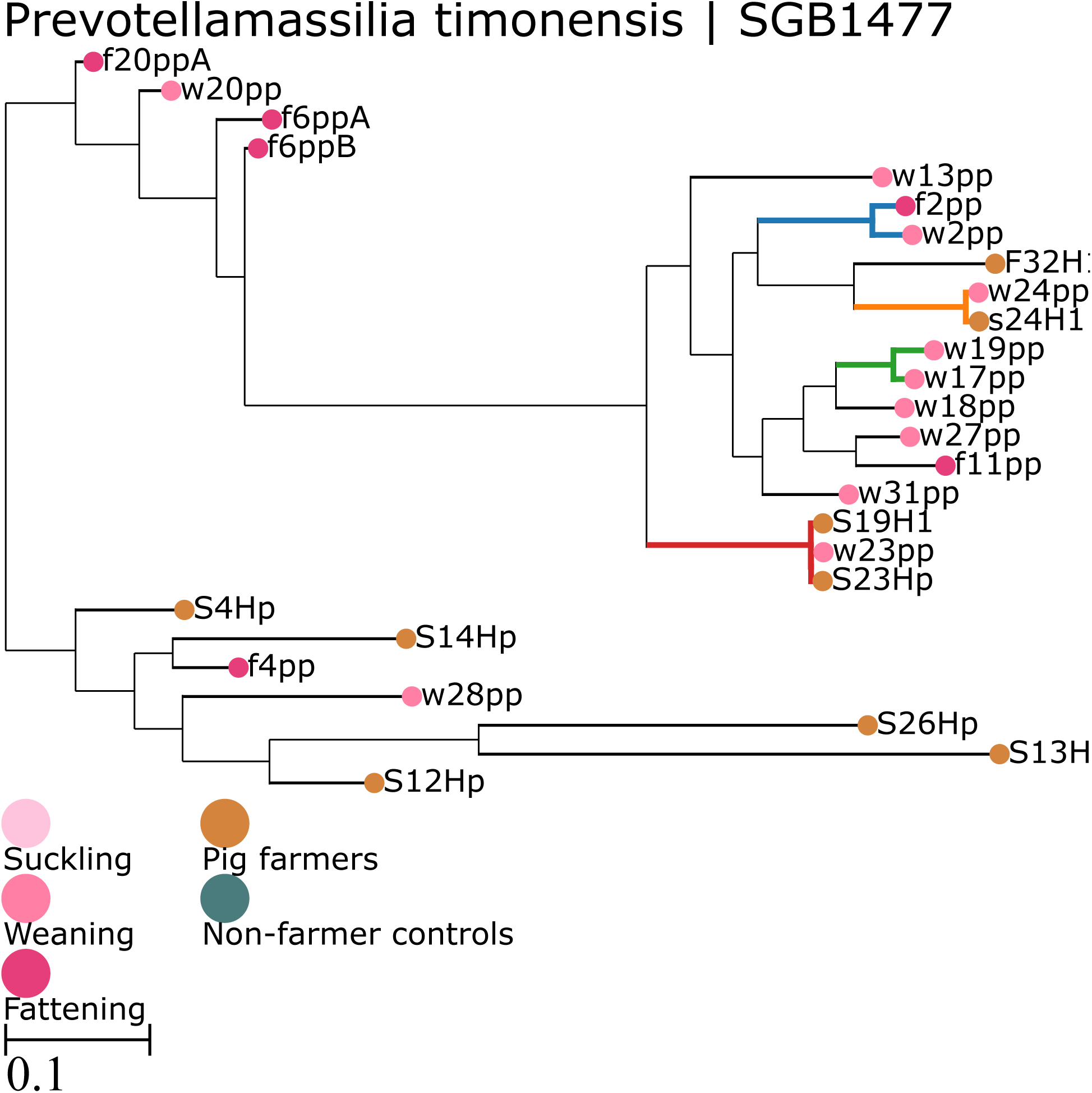
Phylogenetic tree of *Prevotellamassilia timonensis* strains (SGB1477) showing strain sharing between humans and weaning piglets.

**Supplementary Figure S7:**
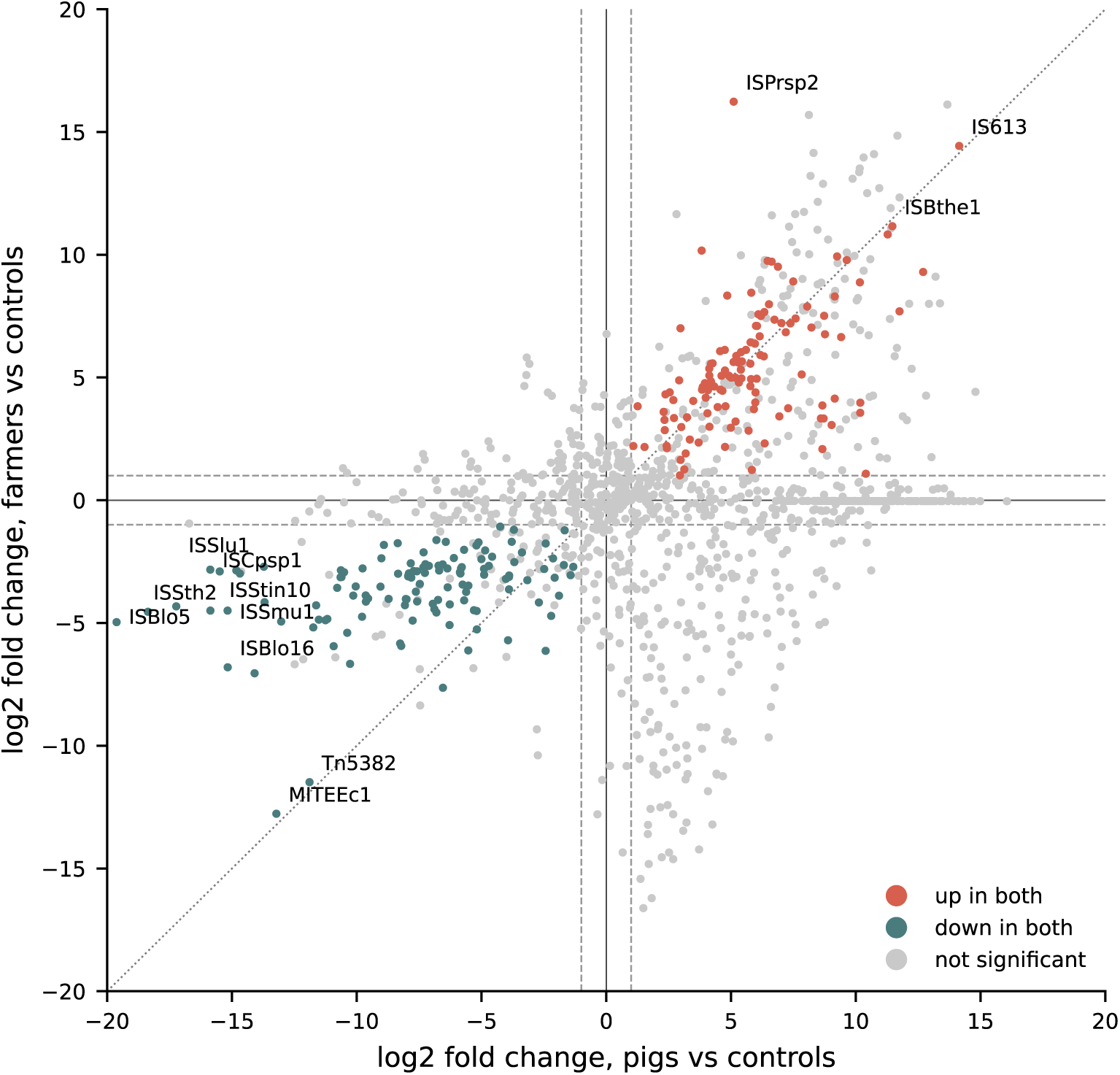
Per-sample differential abundance of all 1,309 MGE elements (after filtering to those detected in 10% of samples) by DESeq2 on element read counts, normalized for sequencing depth. Shown is the log2 fold change for pigs vs controls (x) against farmers vs controls (y); red = significantly up-regulated in both, teal = down-regulated in both, gold = elements of the *Bacteroidetes*-associated IS612/IS613/IS614/IS942/ISBf11/ISBf13/ISPti1/ISMmu1/ISDsu1 families, grey = not significant. The strongest elements are labelled by family.

**Supplementary Figure S8:**
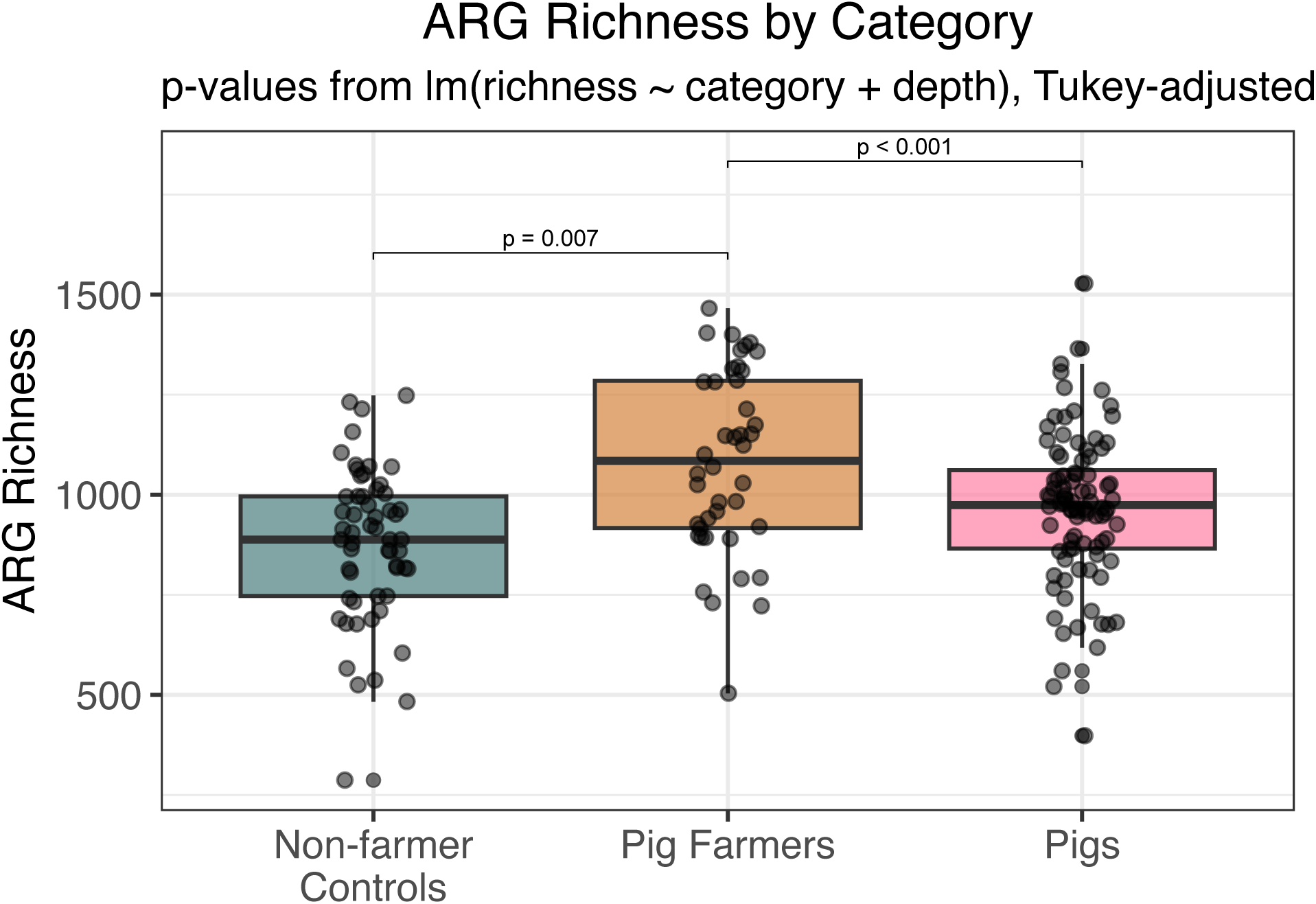
ARG richness by group.

**Supplementary Figure S9:**
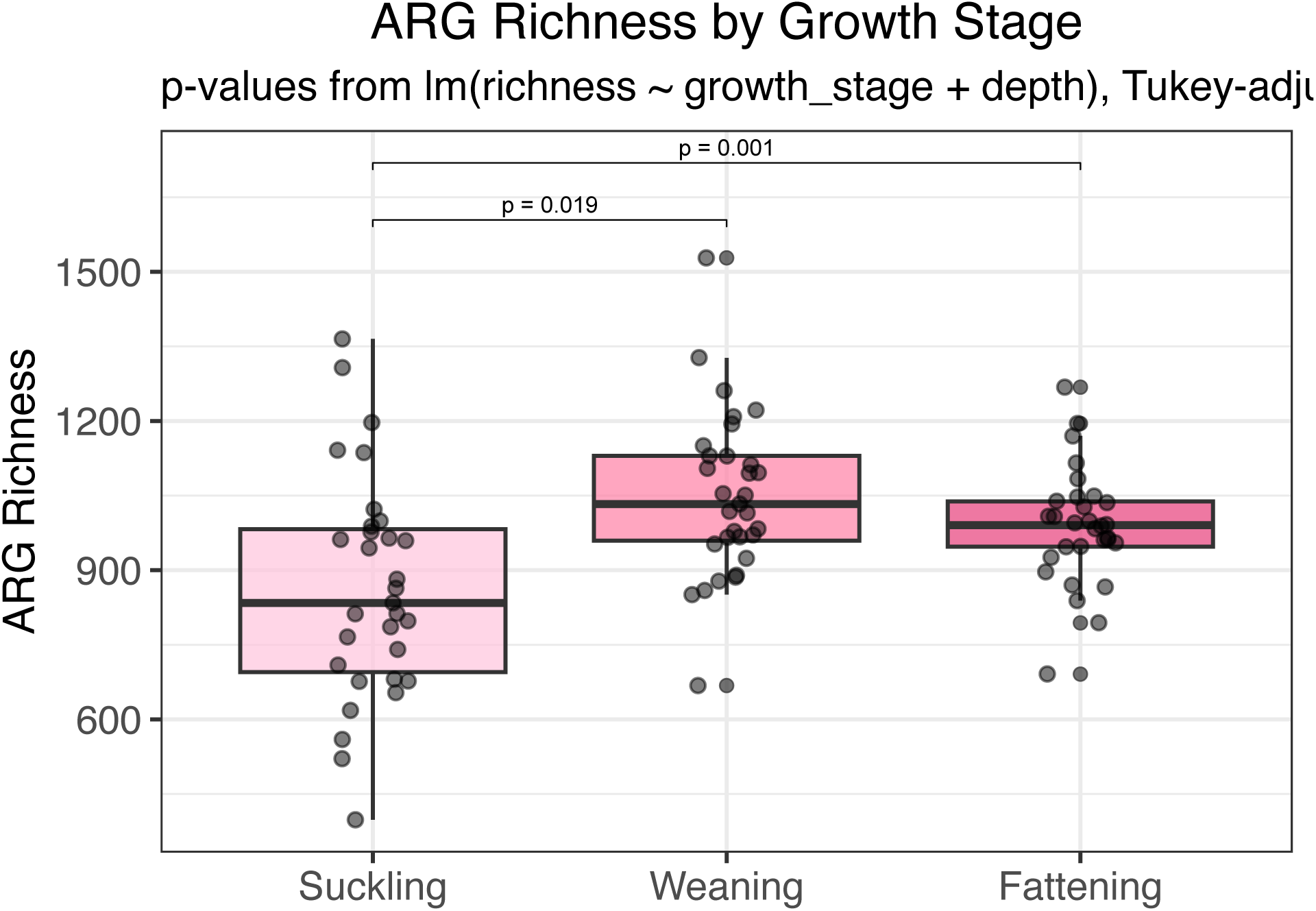
ARG richness by pig growth stage.

**Supplementary Figure S10:**
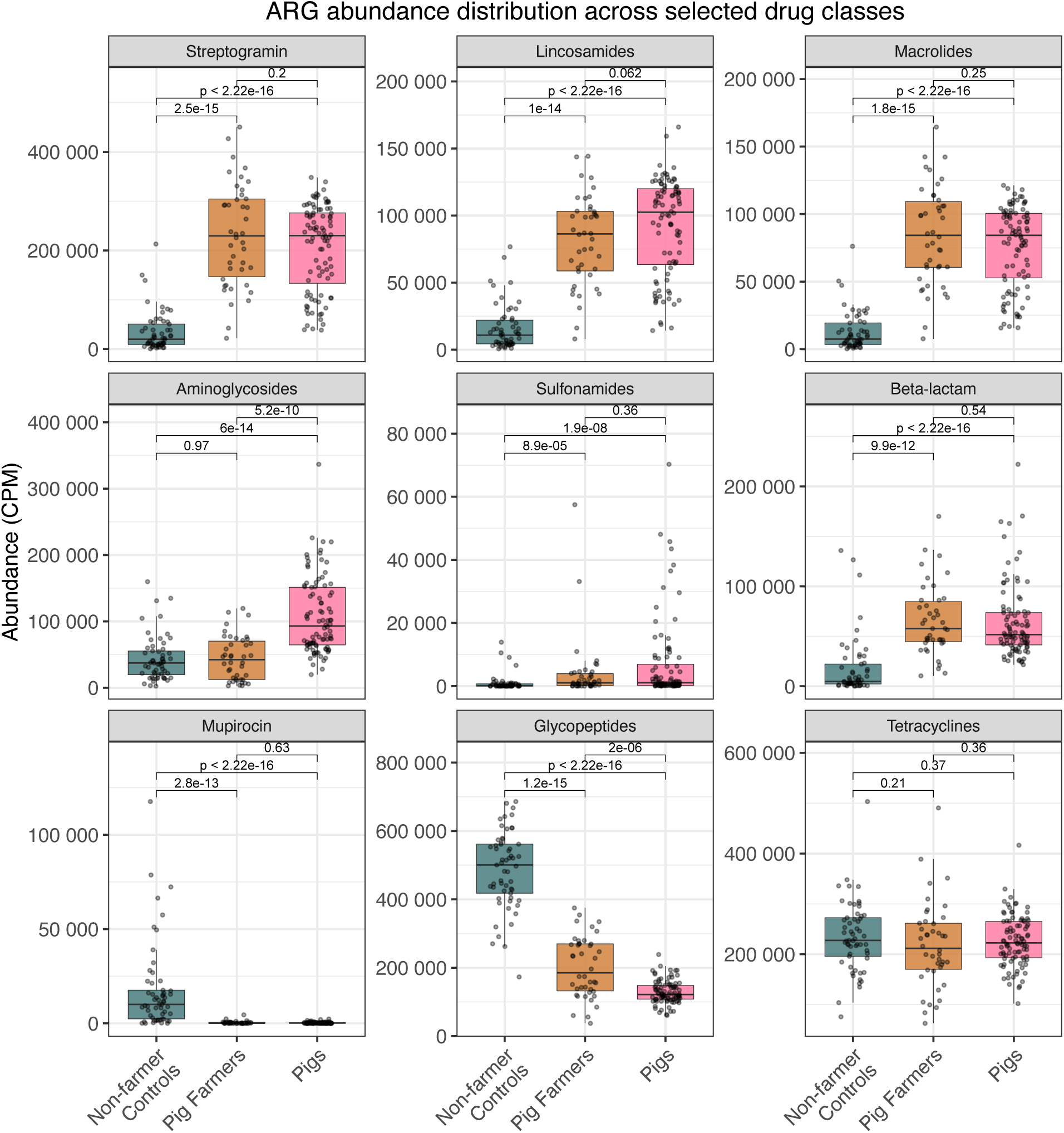
Relative abundance of ARG drug classes by group.

**Supplementary Figure S11:**
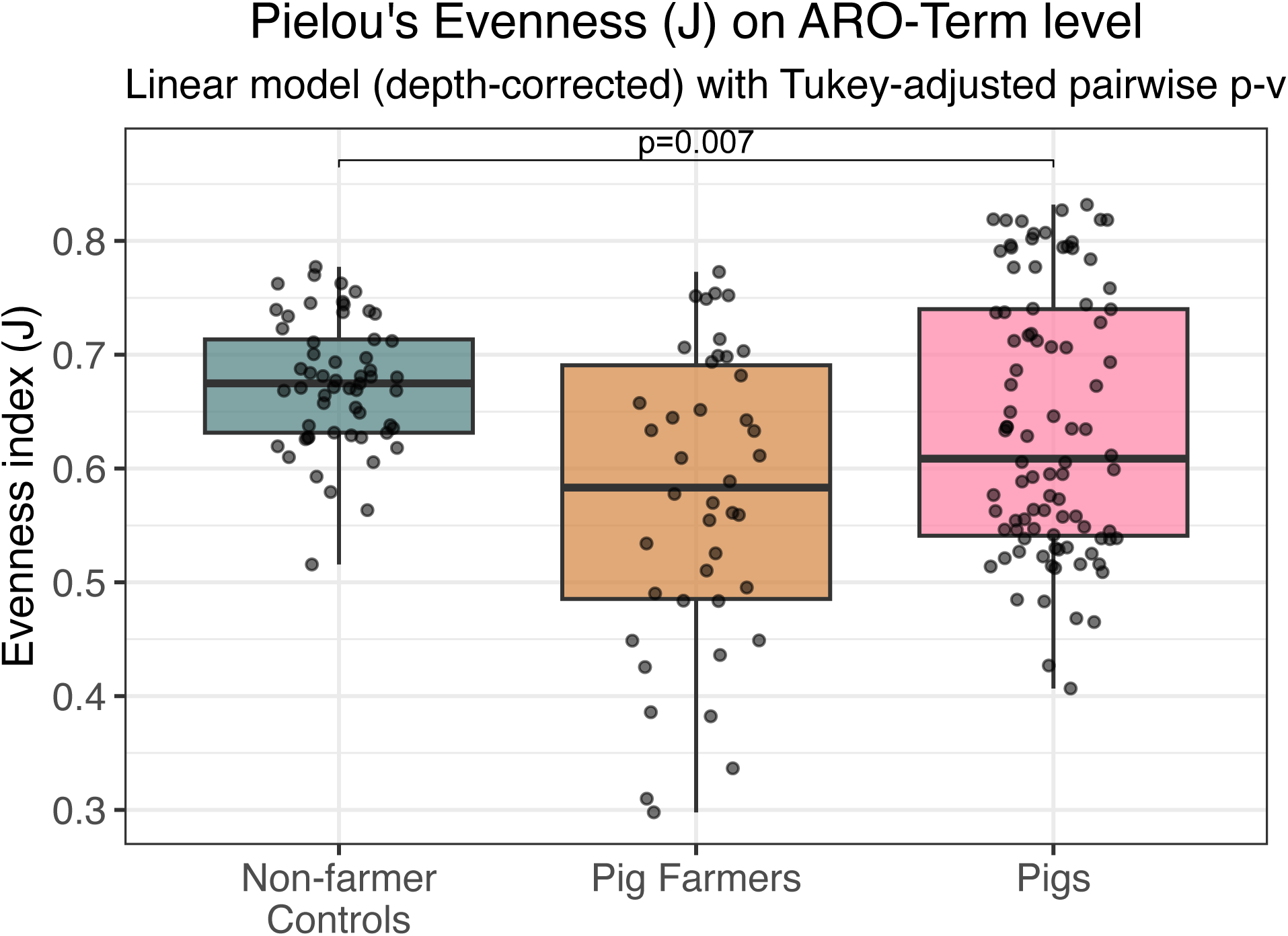
Pielou evenness of ARO terms (depth-corrected).

**Supplementary Figure S12:**
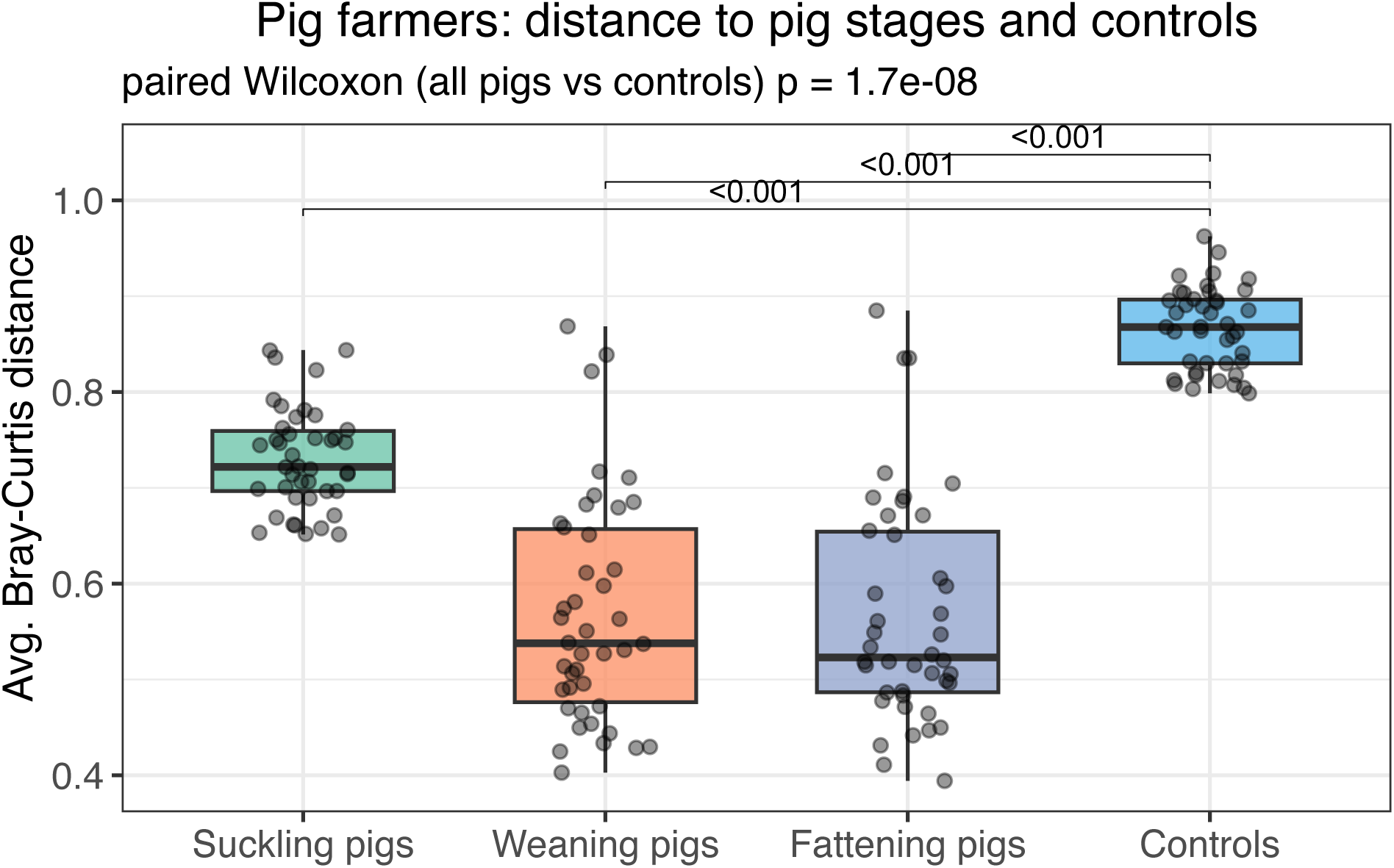
Beta-diversity Bray-Curtis distance from controls and pig farmers to pigs. Pig farmers are significantly closer to pigs (p < 0.001).

**Supplementary Table S1.**
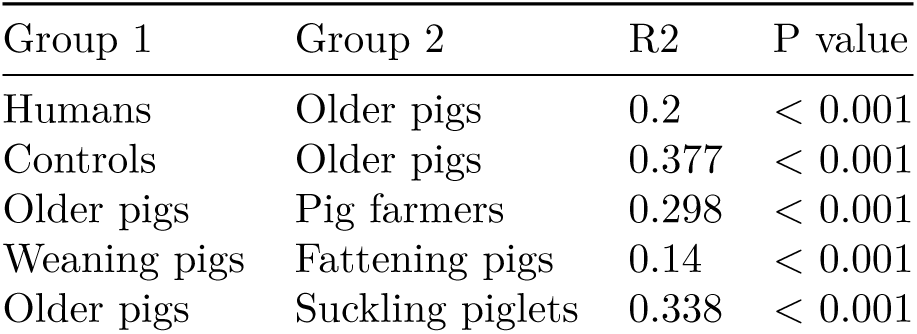

**Supplementary Table S2:**
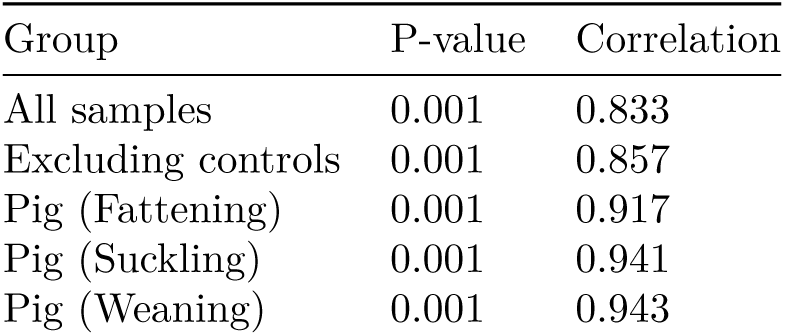
Procrustes analysis: microbiome vs resistome correlation across sample groups.

**Supplementary Table S3:** Administered antibiotics per pig farm.

| Pig Farm ID | Antimicrobials | Timepoint | Drug Class |
| --- | --- | --- | --- |
| 1 | Penicillin | suckling | beta-lactam |
| 2 | Sulfamethoxazol +<br>Trimethoprim | weaning | sulfonamide,<br>diaminopyrimidine |
| 4 | Enrofloxacin | weaning | fluoroquinolone |
| 6 | Marbofloxacin | suckling (sow) | fluoroquinolone |
| 12 | Oxytetracyclin | suckling (sow) | tetracycline |
| 13 | Colistin | weaning | polymyxin |
| 17 | Enrofloxacin | suckling | fluoroquinolone |
| 21 | Sulfamethoxazol +<br>Trimethoprim | suckling | sulfonamide,<br>diaminopyrimidine |
| 23 | Lincospectin | suckling | lincosamide |
| 28 | Sulfamethoxazol +<br>Trimethoprim | suckling | sulfonamide,<br>diaminopyrimidine |
| 30 | Amoxicillin | suckling | beta-lactam |
| 32 | Amoxicillin + Clavulanic<br>acid + Oxytetracyclin | suckling | beta-lactam,<br>tetracycline |

**Supplementary Table S4:** MGE linkage was defined as an MGE predicted by MEFinder within 1 kb of the ARG on the same assembled contig. Only ARGs with detectable MGE association are shown; the remaining ARGs elevated in both pigs and farmers (*cfxA4*, *cfxA2*, *dfrD*, *cfrE*, *aadA9*, *ANT(3 )-IIa*, *APH(2 )-IIa*, *AAC(6 )-Im*, *lsaE*, *mel*, *tet(O/W)*, *OXA-193*, *ermQ*, *tet(W/32/O)*, *vanW*, *nimB*) had no appreciable MGE linkage ( 2% of loci).

| Gene | Family | Loci (n) | MGE-linked loci (n) | % MGE-linked | MGE types |
| --- | --- | --- | --- | --- | --- |
| *tet(X)* | tetracycline<br>inactivation | 81 | 64 | 79% | IS4351;<br>Tn4351 |
| *ermF* | Erm 23S<br>rRNA<br>methyl-<br>transferase | 94 | 43 | 46% | ISBf11;<br>Tn4351 |
| *dfrA1* | dihydrofolate<br>reductase | 26 | 4 | 15% | Tn2; Tn7 |
| *sul2* | sulfonamide | 70 | 10 | 14% | ISBbi1; ISVsa3 |
| *estT* | macrolide<br>esterase | 9 | 1 | 11% | IS4351 |
| *aadA* | ANT(3 ) | 57 | 4 | 7% | Tn7; IS6100 |

**Supplementary Table S5:** The five strain-sharing events between pig farmers and fattening pigs (nGD < 0.1). * Same farm but during suckling and weaning stage.

| Farmer | Fattening<br>pig | Farm relationship | Species | Family | nGD |
| --- | --- | --- | --- | --- | --- |
| S1Hp | f1pp | Same Farm* | SGB5809 | Veillonellaceae | 0.050 |
| S1Hp | f18pp | Different farm | SGB6293 | FGB1765 | 0.100 |
| S30H1 | f10pp | Different farm | SGB70108 | Prevotellaceae | 0.007 |
| F16H1 | f16ppA | Same farm | *A. ermentans* | Acidaminococcaceae | 0.062 |
| s24H1 | f24pp | Same Farm* | *Flintibacter* SGB15145 | *Eubacteriales* unclassified | 0.076 |

**Supplementary Table S6:**
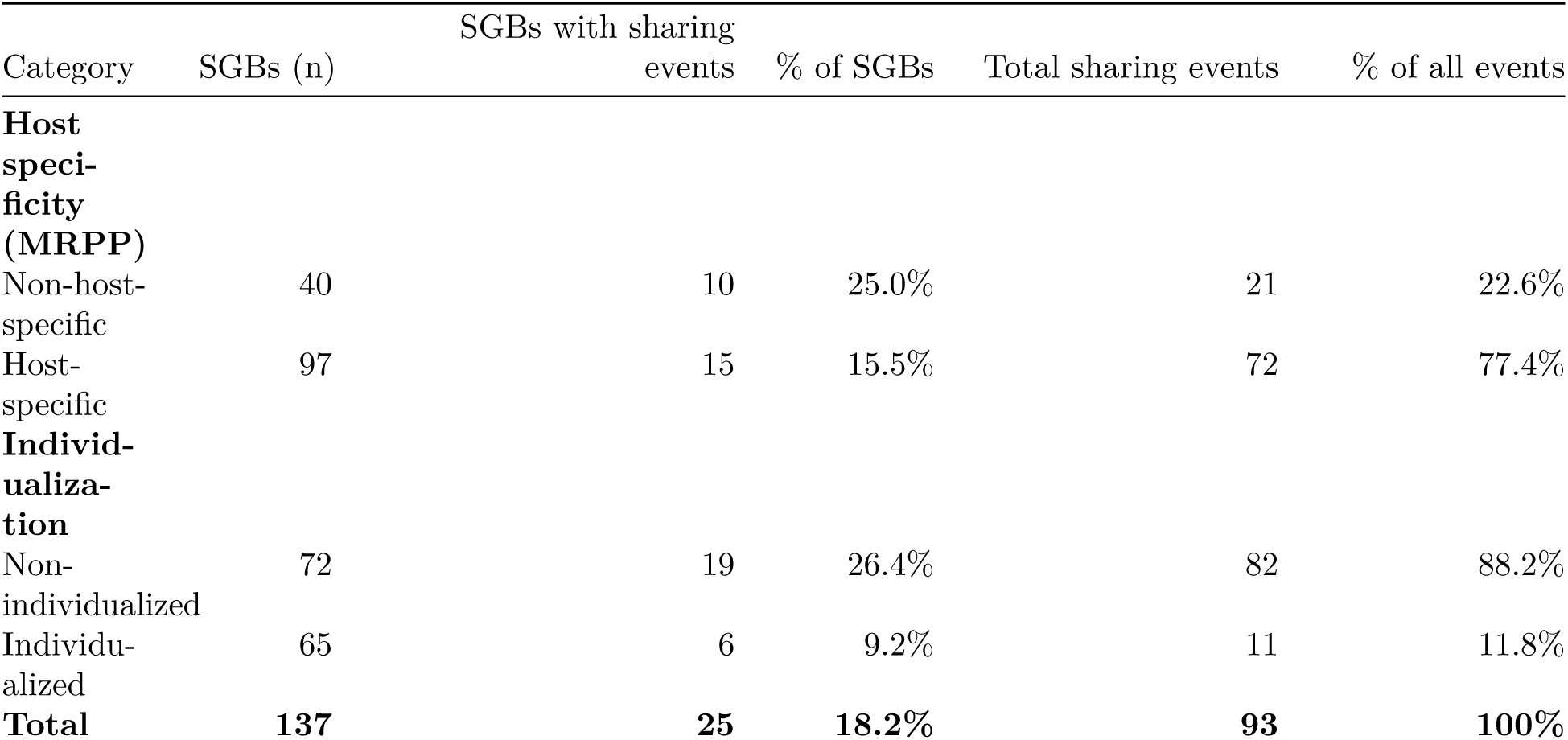
Strain sharing events (farmer–pig pairs, nGD < 0.1) tabulated by host-specificity (MRPP) and individualization score category.

